# A Reactive Inelasticity Theoretical Framework for modeling Viscoelasticity, Plastic Deformation, and damage in Soft Tissue

**DOI:** 10.1101/247510

**Authors:** Babak N. Safa, Michael H. Santare, Dawn M. Elliott

**Author notes:** This document is a collaborative effort. Corresponding author: *Email address:* (Dawn M. Elliott). 150 Academy Street, 161 Colburn Lab, Newark, DE 19716.

## Abstract

Soft tissues are biopolymeric materials, primarily made of collagen and water. These tissues have non-linear, anisotropic, and inelastic mechanical behaviors that are often categorized into viscoelastic behavior, plastic deformation, and damage. While tissue’s elastic and viscoelastic mechanical properties have been measured for decades, there is no comprehensive theoretical framework for modeling inelastic behaviors of these tissues that is based on their structure. To model the three major inelastic mechanical behaviors of soft tissue we formulated a structurally inspired continuum mechanics framework based on the energy of molecular bonds that break and reform in response to external loading (reactive bonds). In this framework, we employed the theory of internal state variables and kinetics of molecular bonds. The number fraction of bonds, their reference deformation gradient, and damage parameter were used as internal state variables that allowed for consistent modeling of all three of the inelastic behaviors of tissue by using the same sets of constitutive relations. Several numerical examples are provided that address practical problems in tissue mechanics, including the difference between plastic deformation and damage. This model can be used to identify relationships between tissue’s mechanical response to external loading and its biopolymeric structure.

## 1. Introduction

Soft tissues such as tendon, meniscus, intervertebral disc, etc., are biopolymeric material that have non-linear, anisotropic, and inelastic mechanical behaviors. An inelastic behavior, contrary to an elastic behavior, is one that dissipates energy in a non-recoverable fashion. Three major inelastic behaviors that are experimentally observed in soft tissues are: viscoelastic behavior (Woo et al., 1980; Huang et al., 2001; Connizzo & Grodzinsky, 2017), plastic deformation (Maher et al., 2012; Caro-Bretelle et al., 2015), and damage (Natali et al., 2005; Von Forell & Bowden, 2014). Understanding the underlying mechanisms of these behaviors is essential for elucidating the relationships among the mechanical loading, pathological conditions (e.g., tendinopathy, meniscus rupture, and disc herniation), and tissue’s structure.

Soft tissues have a hierarchical fibrous structure that is predominantly composed of collagen and water, where the underlying microstructural organization is responsible for the variations in tissue’s mechanical properties. Inelastic mechanical responses can occur with loading, resulting in altered microstructure and mechanical properties, and even loss of function and rupture. While tissue’s elastic and viscoelastic behaviors have been studied for decades, there is no comprehensive framework for studying inelastic behaviors of tissue in a structural context; perhaps due to the complex nature of experimental measurement of the inelastic behaviors that occur simultaneously and can have overlapping effects (e.g., plastic deformation and damage). Thus, a theoretical framework is necessary to study these behaviors that uniquely identifies characteristics of different inelastic behaviors and their underlying mechanisms based on tissue structure.

From a theoretical point of view, an inelastic behavior is a path-dependent and irreversible thermodynamic process that may be modeled using the theory of internal state variables (ISV). ISVs are mathematical variables that are included in the definition of the state of material in addition to the ‘state variables’ (e.g., deformation measures, temperature, etc.) These variables may be used to account for the inelastic mechanical response that is non-recoverable (Coleman & Gurtin, 1967; Horstemeyer & Bammann, 2010). Various phenomenological ISV expressions have been used to model tissue’s mechanical behavior. For instance, decomposition of strain (Reese & Govindjee, 1998) and stress (Holzapfel & Simo, 1996) tensors into elastic and inelastic components is a common practice to model viscoelasticity, where the inelastic components are hidden internal variables that correspond to the non-equilibrium part of the mechanical response. For plastic deformation (permanent set) in tissue (Zhang & Sacks, 2017) used a structural approach to model this inelastic behavior in cyclic loading by proposing a shift in reference configuration of exogenously crosslinked matrix of tissue. In another study, to model softening based on permanent set and damage two weight factors were used based on alterations to the fibrous structure of tissue due to mechanical loading (Peña, 2011, 2014). A similar inelastic variable set was used to model permanent deformation and damage that evolve according to the history of maximum energy of fiber families (Fereidoonnezhad et al., 2016) that is based on the theory of *pseudo-elasticity* (Weisbecker et al., 2012; Dorfmann & Ogden, 2004). Damage, perhaps the most common ISV for modeling tissue inelasticity (Li, 2016), is often used to model stress-softening or Mullin’s effect (Mullins, 1969; Diani et al., 2009). In tissue, damage is usually modeled according to the number of “broken” fibers (Natali et al., 2005; Schmidt et al., 2014; Alastrué et al., 2007). This formulation is similar to the definition of the damage parameter in classic continuum damage mechanics framework that is based on occurrence of micro-voids in engineering material such as metals and concrete (Kachanov, 1968; Lemaitre, 1984). Despite nearly four decades of successful use of phenomenological approaches to study tissue’s inelastic behaviors, their applicability is limited by the focus of individual models on a subset of inelastic behaviors, and in particular for ISV approaches, there is often not a clear physical interpretation of parameters.

In this study, we used the theory of ISVs and the kinetics of molecular bonds to develop a structurally inspired continuum mechanics framework that addresses the inelastic behaviors of tissue. In a few words, we define reactive bonds as those which break and reform in response to mechanical loading; this process results in dissipative changes to the stored energy of the material and inelastic mechanical behaviors. We selected the number fraction of molecular bonds (*w*^α^), their reference deformation gradient (**Π**^α^), and damage parameter (*D*) as ISVs. This framework allows for modeling of all three of the aforementioned inelastic behaviors, viscoelastic behavior, plastic deformation, and damage, using the same set of constitutive relations.

Kinetics of molecular bonds has been previously used by Tobolsky and co-workers to model stress-relaxation of polymers based on the energy of a network of molecules with transient cross-links. This is also commonly referred to as the *two-network model* (Green & Tobolsky, 1946; Tobolsky & Andrews, 1945). In a similar way, permanent set has been modeled using the idea of kinetics of breaking and reforming bonds (Andrews et al., 1946; Rajagopal & Wineman, 1992), where often the idea of material with multiple natural configurations is employed in a thermomechanical framework (Rajagopal & Srinivasa, 2004; Muliana et al., 2016). This framework is widely used for studying mechanical behaviors of engineering polymers (Scott & Stein, 1953; Demirkoparan et al., 2009; Wineman, 2009; Meng et al., 2016) with various physical interpretations based on the molecular structure of polymers (Diani et al., 2009). For tissue mechanics application, a similar idea was recently adapted to model damage of cartilaginous constructs that addresses damage as the fraction of failed bonds, which is an observable variable and can be experimentally measured (Nims et al., 2016). Reactive viscoelasticity was introduced by Ateshian to model viscoelasticity of a constrained solid mixture (Ateshian, 2015; Nims & Ateshian, 2017). In that model, two separate bond types were introduced: weak and strong, to model the transient and equilibrium mechanical response, respectively. In the current work, we generalize that concept and show that these bonds are not separate, and strong bonds are a special case of weak bonds. Additionally, by employing this generalization and taking inspiration from prior work in polymer mechanics, the sliding bond type was introduced that can instantaneously break and reform into a stressed-configuration, which results in plastic deformation. Given the structural basis of this framework and its consistency in addressing several inelastic behaviors, reactive inelasticity is well suited for studying the underlying mechanisms of inelasticity in tissue. Thus, the objective of this study was to comprehensively address tissue inelasticity by developing a structurally inspired continuum mechanics framework, reactive inelasticity, based on the energy of reactive molecular bonds.

This paper is organized as follows: in Section 2 we formulate the theory of reactive inelasticity by defining the free energy of a reactive body with generic molecular bond types, where the concepts of intrinsic hyperelasticity and generalized bond kinetics are formulated. Further, we introduce special bond types suited for different mechanical behaviors: formative bonds for a viscoelastic behavior, permanent bonds for hyperelastic behavior, and sliding bonds for plastic deformation. Damage is added to the formulation by allowing for the reduction of the number fraction of load bearing bonds. To demonstrate the key features of this framework, in Section 3 we provide several numerical examples of the mechanical behavior of single bond types and their combinations. Finally, in Section 4, comparisons to the other existing inelasticity models for tissue, and physical interpretation of bonds are discussed.

## 2. Reactive inelasticity

In the following, we will describe the mechanics of a reactive material. In brief, a reactive material is made of a combination of different bond types each with different characteristics. Each bond type is associated with a certain mechanical behavior (e.g., formative bonds with viscoelastic behavior). Bonds break and reform (i.e., react) when subjected to an external loading and this process initiates new generations. Generations are molecular bonds of certain bond type that were initiated at the same time and, thus, have a mutual reference configuration. By using the kinetics rate of bond breakage and formation, the states of energy and stress are determined. Damage is introduced to all of the bond types by reducing the fraction of active bonds in the material, and finally the consequences of the second law of thermodynamics are discussed.

### 2.1. Kinematics and relative deformation

Consider a solid body made of a set of material points that are energetically constrained with different types of molecular bonds (indicated by a subscript *γ*). These bonds deform in a smooth path 
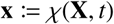
, where **x** stands for the current configuration of bonds and **X** is their master reference configuration. Hence, the deformation gradient tensor is defined as 
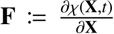
 and the right Cauchy-Green deformation tensor as **C**:= **F**^*T*^ **F**. Each bond type may have different generations (indicated by superscript *α*), and despite their mutual path of deformation and master reference configuration, every generation has a different reference configuration 
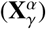
 which is a function of the conditions at the time of formation of the bonds (Fig. 1). By using the chain rule of differentiation, the relative deformation gradient tensor for each generation 
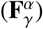
 is 

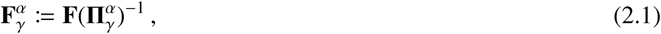

where 
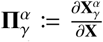
 is the reference deformation gradient tensor of generation *α* of bond type *γ*. This procedure results in *multiplicative decomposition* of deformation (Lee, 1969; Simo, 1988; Lubarda, 2004). As a result, the relative right Cauchy-Green deformation tensor 
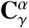
 is

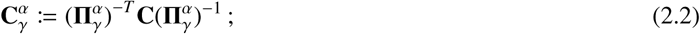

note that 
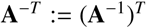
.

### 2.2. Free energy and intrinsic hyperelasticity

The free energy density of the system is the combination of the energy density of all the bond types

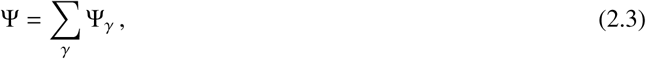

where Ψ is the Helmholtz free energy density (per volume). The energy density of each bond type is the sum of the energy from each of its generations

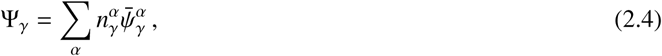

where 
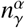
is the number density, and 
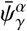
 is the average energy of bonds from generation *α* of bond type *γ*. Alternatively, by defining the number fraction of each generation as

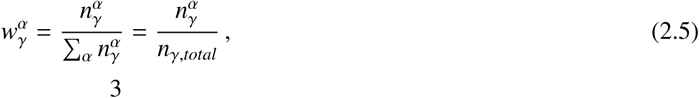

**Figure 1.**
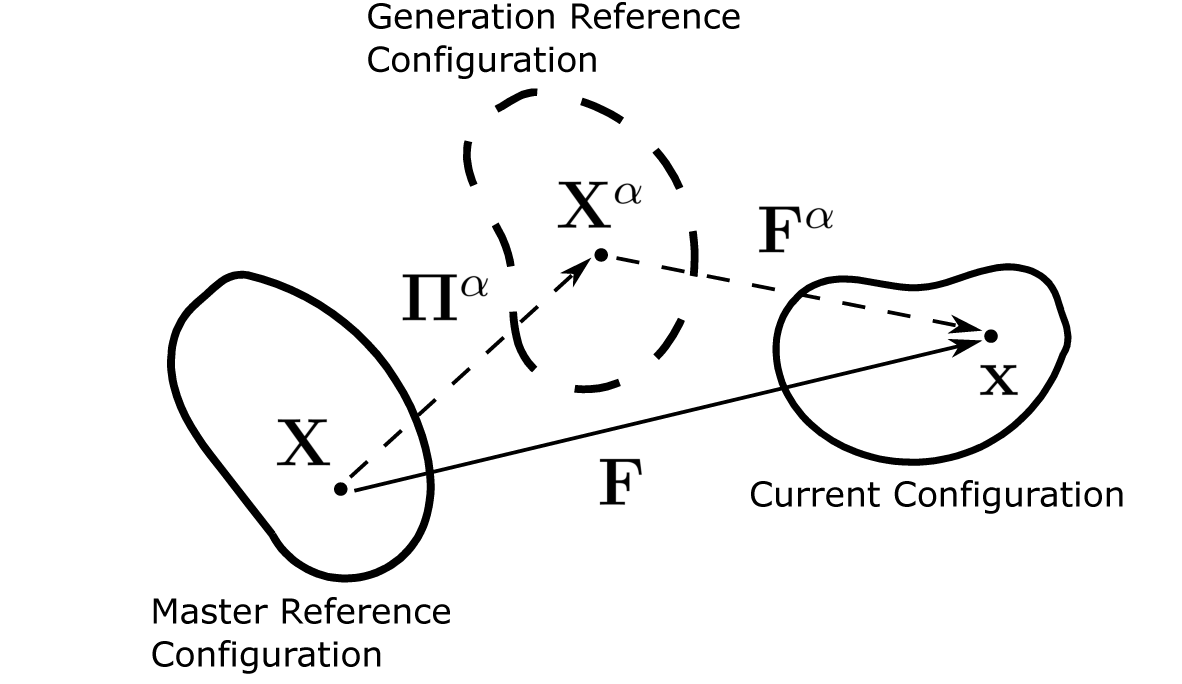
Reference configuration of generations. In a body with the master reference configuration **X** for generation the relative deformation gradient tensor (**F**^*α*^) is defined using the reference configuration (**X**^*α*^).

where *n_*γ*, total_* is the number density of a bond type in the natural state of the material, Eq. (2.4) can be re-written as

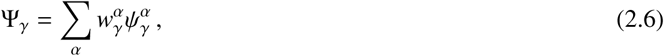

where

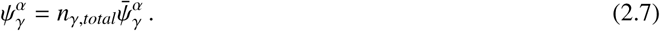

Here, 
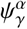
 denotes the overall energy density of a generation, which, in general, can be a function of deformation, temperature and other state variables. However, in this formulation we consider a unique deformation dependence that implies that generations are *intrinsically hyperelastic*. That is,

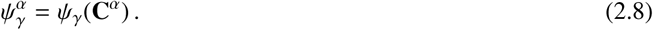

The above equation also denotes that the intrinsic hyperelasticity relation is the same for all of the generations of one bond type, and thus is a characteristic of that type of bonds. Therefore, Eq. (2.6) reads as

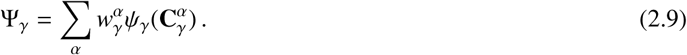

Note that for a bond type with no damage

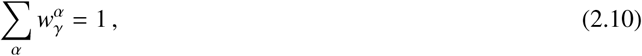

which indicates that the energy density of each bond type (Eq. (2.9)) is a weighted sum of the energy density of all its generations. Thus, the overall free energy density of the material can be written in the following form

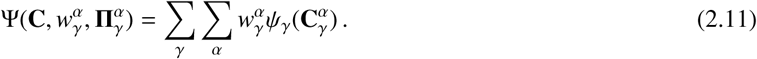

Here, deformation (represented by **C**) is the only “state variable” and the number fraction of generations of bond types 
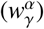
 as well as their reference deformation gradient 
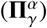
 are the “internal state variables (ISV)”. To simplify the notation without losing generality, we will continue the discussion for one type of bond, where the total response for a mixture of multiple bond types should be taken as a summation over all existing bond types (such as in Eq. (2.11)).

The second Piola-Kirchhoff stress tensor is written as (Coleman & Gurtin, 1967)

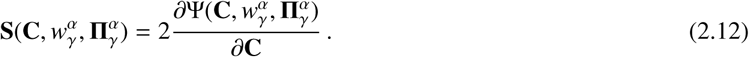

Also, by using Eq. (2.9) for a reactive material, Eq. (2.12) can be re-written as (see Appendix A)

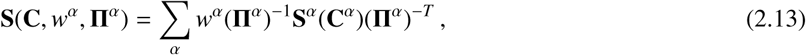

where we also adopted the following definition for stress associated with each generation:

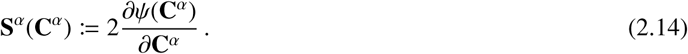

To enforce coordinate-invariance, the state of stress equivalently is determined using the invariants of C^*α*^:

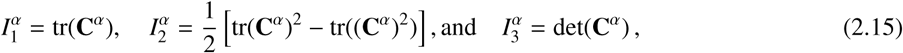

where tr(.) and det(.) stand for trace and determinant of a second order tensor, respectively. In general, the given set of invariants suffices to define the state of deformation; however, it is customary to add additional *pseudo invariants* for anisotropic material (e.g., tendon, and ligament). A common pseudo invariant for transversely isotropic material is defined as (Spencer, 1984; Cortes & Elliott, 2014)

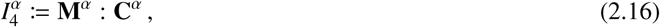

In this equation, **M**^*α*^ is a second-order structure tensor defined as

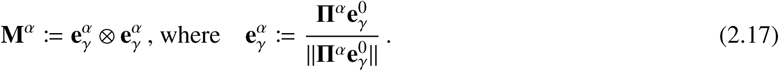

Here, 
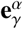
 is a unit vector on the dominant fiber direction in the generation *α* reference configuration. Therefore, the second Piola-Kirchho stress (**S**) in terms of deformation invariants reads as

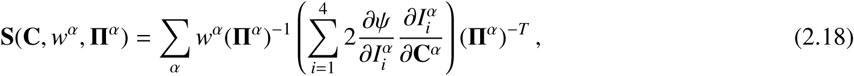

where the derivatives of the deformation invariants are

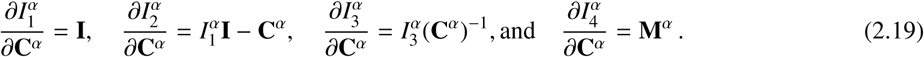

The Cauchy stress (**T**) is then obtained from the second Piola-Kirchhostress (Eq. (2.18)) as

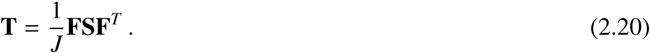

By using the relative deformation gradient tensor, defined earlier in Eq. (2.1), the Cauchy stress tensor for each bond type is

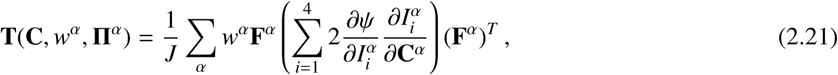

and by defining the contribution of each generation (**T**^*α*^) as

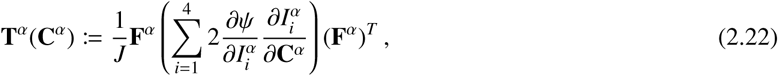

the Cauchy stress tensor for a specific bond type reads as

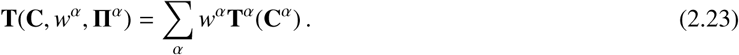

Similar to energy, the above equation indicates that the Cauchy stress of a bond type is the weighted sum of the contributions from all the generations of that bond type.

So far, we have described the fundamental formulation for states of energy and stress in terms of state variables (**C**,*w*^*α*^,**Π**^*α*^). In the following sections, we will describe the internal state variables and how they evolve to describe the states of energy and stress in a reactive material.

### 2.3. Reactive bond kinetics

In general, the reactive bonds break and reform (react) when subjected to loading. This process produces the evolution of the states of energy and stress (Eqs. (2.11) and (2.23)). As an example, for a system with one type of bonds, after a unit step deformation the bonds start breaking and their number fraction (*w*^0^) starts to decrease (Fig. 2). Simultaneously, the broken bonds reform to a new state, initiating a new generation, where the number fraction of this new generation (*w*^1^) increases over time. In the absence of damage, according to Eqs. (2.10) the sum of the number fraction of the breaking and reforming bonds is one (i.e., *w*^0^(*t*) + *w*^1^(*t*) = 1). As a result, the free energy for a step deformation is

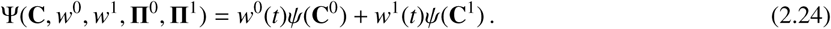

If the bonds reform in a stress-free fashion (**C**^1^ = **I**), without further loading all of the energy would be dissipated. That is,

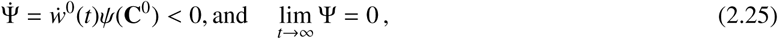

where 
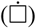
 stands for time derivative. To determine the rate of breakage of bonds, a kinetics rate equation is used. The general form of the kinetics rate equation may be written as

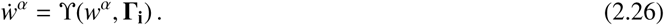

In this equation, 
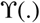 is a negative-valued function of the available reactants (breaking bonds of *w*^*α*^) and other state variables such as relative deformation (**Γ** = **F**^*α*^) (Ateshian, 2015), temperature (Γ = *θ*) (Wineman, 2009), or for polymeric macro molecules, geometrical characteristics of bonds such as the end-to-end distance and directional angles (Meng et al., 2016; Tanaka & Edwards, 1992). Despite features that can be added to the rate equation, there are general conditions that need to be satisfied. First, any rate equation should not result in a negative number fraction for any bond generation. Secondly, it should induce a breaking bond number fraction that asymptotically decays to zero in the limit of infinite time. In this paper, we used the first-order relation, which is the simplest form of a kinetics rate equation satisfying the above conditions:

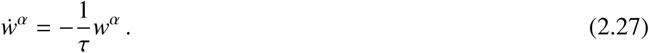

In the above equation *τ* is the time constant of reaction that controls the rate of breakage and reformation. When multiple consecutive steps of deformation occur at 
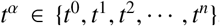
, the number fraction resulting from each deformation for this first-order system is (Fig. 3A) (Ateshian, 2015)

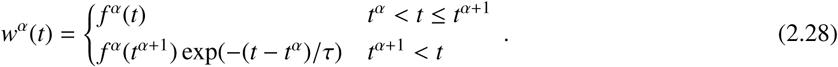

In the above relation 
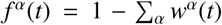
 according to Eq. (2.10) that describes the reformation of bonds. The dependence of the rate of breakage on the number fraction of breaking bonds results in an asymptotic exponential decay of bonds. Note that Eq. (2.8) can only be used for first-order kinetics with non-varying time-constant, and for a general rate equation, the number fraction of bonds should be directly calculated from Eq. (2.10) and (2.6).

### 2.4. Bond types

The state of energy (Eq. (2.11)) for a given bond type depends on the number fraction (*w*^*α*^) of bonds and their reference deformation gradient (**Π**^*α*^). To characterize the evolution of these internal state variables we define three types of bonds: (1) formative, (2) permanent, and (3) sliding to address viscoelasticity, hyperelasticity, and plastic deformation, respectively.

#### 2.4.1. Formative bonds

These bonds have a time constant that is on the order of the characteristic time (*t_c_*) of the experimental observation. That is, *τ*_*f*_ / *t_c_* ≈ 1, where *γ* = *f* stands for formative bond type (Fig. 3A). The reference deformation gradient of the new generations of bonds is defined at the current configuration at the time the bonds were initiated (Fig. 3D)

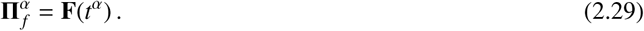

**Figure 2.**
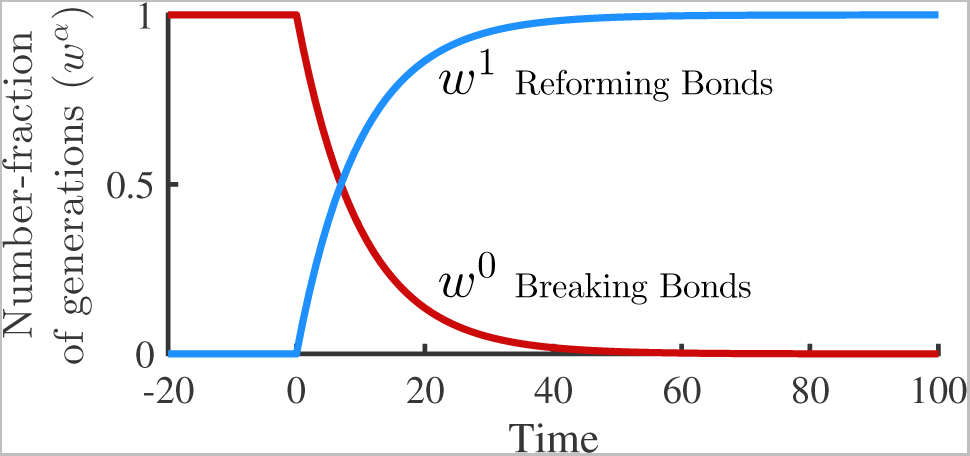
Example of bonds with first-order rate equation. After a step deformation at *t* = 0, the bonds start to break (*w*^0^), and simultaneously reform to a new state (*w*^1^). With no further loading, all the bonds eventually break and reform to the new configuration as *t* → ∞ (in this example *τ* = 10).

Equivalently, for two consecutive generations

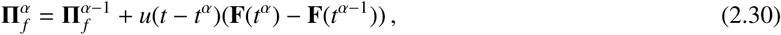

where *u*(.) is the unit step function. In this case, during a sustained deformation the energy and stress of the system asymptotically decays as the bonds break, resulting in a fluid-like behavior; a transient viscoelastic mechanical response with zero equilibrium stress.

#### 2.4.2. Permanent bonds

Permanent bonds represent a special case of kinetics rate, where the rate of breakage of bonds is extremely slow 
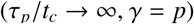
 (Fig. 3B). As a result, only one generation of bonds exists 
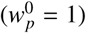
 and the reference deformation gradient of that generation is (Fig. 3E)

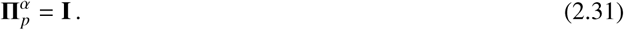

Since the bonds are intrinsically hyperelastic (Eqs. (2.8) and (2.14)), the overall behavior of permanent bonds is hyperelastic, where the states of energy and stress are independent of the history of deformation.

#### 2.4.3. Sliding bonds

Sliding bonds represent another special case, where the rate of breakage and reformation of bonds is extremely fast compared to the characteristic time 
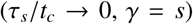
 (Fig. 3E). In this case, if the bonds could reform to the current configuration, all the energy would immediately be dissipated. If the newly formed bonds have a different reference deformation gradient 
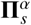
 than the current configuration, they slide (Fig. 3F). This implies that the bonds are reforming in a loaded state, and only part of the energy is used for the sliding process and the rest is stored. Since 
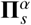
 is an internal state variable, by selecting appropriate constitutive relations the sliding process can be made to produce plastic deformation.

The evolution of 
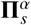
 is governed by a set of constitutive relations. Following from the classical theory of plasticity (Simo & Hughes, 1998; Khan & Huang, 1995) we formulated the evolution of the sliding reference deformation gradient using a sliding condition and sliding rule in a rate-independent formulation. Similar to Eq. (2.30) the sliding process is incremental from one generation to the next, i.e.,

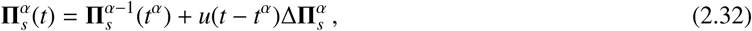

where 
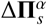
 is an incremental infinitesimal change to the reference deformation gradient due to sliding. Equation (2.32) can be also written as 
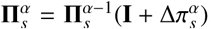
, where 
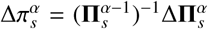
; then it follows that the infinitesimal sliding strain is calculated as 
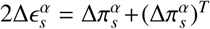
, which provides an explicit relation between the infinitesimal sliding strain and the incremental change to the reference deformation gradient of a generation. Note that the zeroth generation’s the identity tensor (**I** = **e**_*i*_ ⊗ **e**_*i*_). Although the change is infinitesimal, it does not imply that 
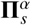
 is infinitesimal, nor that the incremental nature of the sliding process goes away, no matter how small the sliding would be.

**Figure 3.**
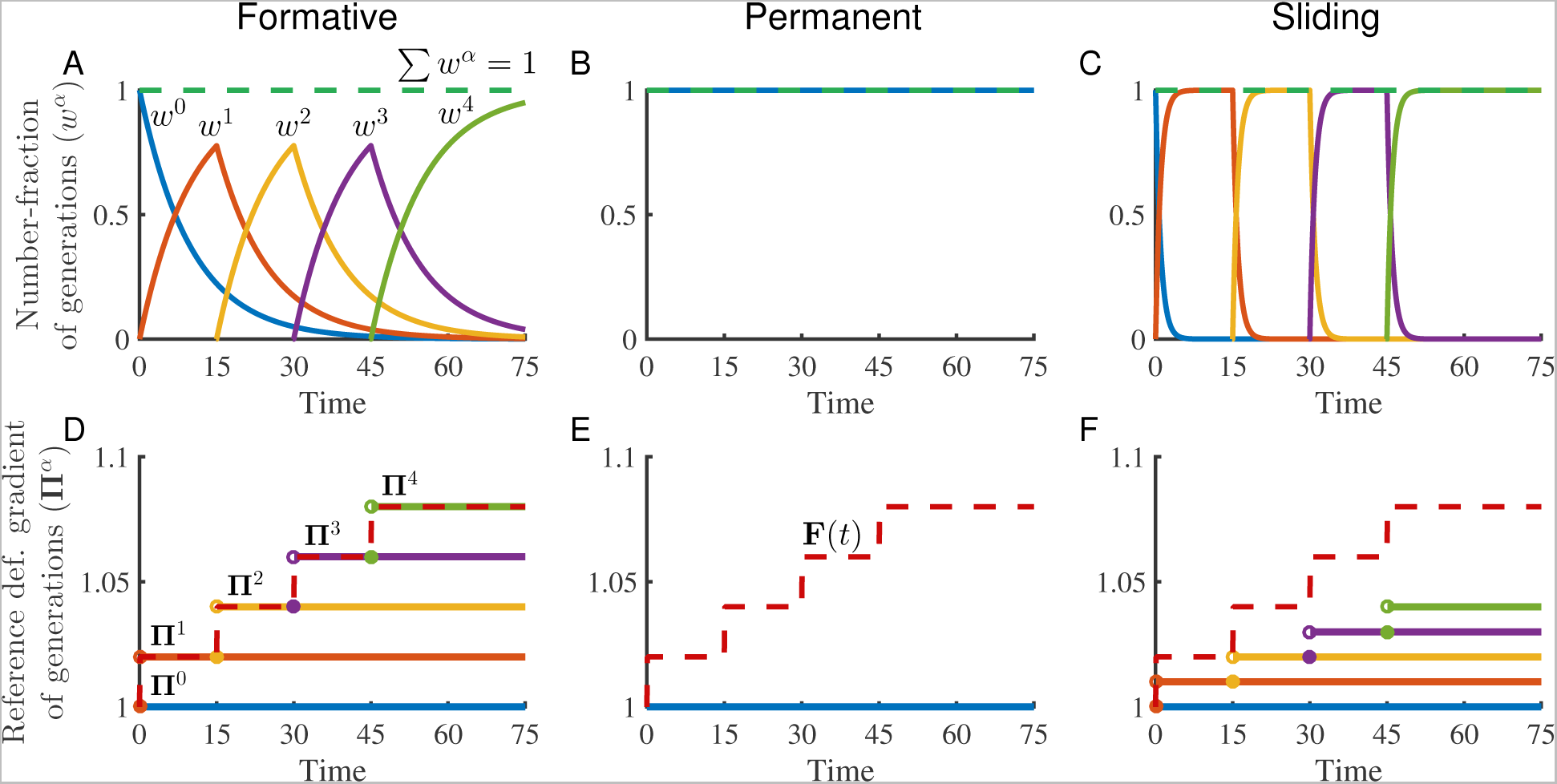
Consecutive step deformations and multiple generations. (A-C) Number fraction of multiple generations initiated at four consecutive step deformations at *t* = 0, 15, 30, 45 with a first-order rate equation, and (D-F) their corresponding reference deformation gradients. (A,D) Formative bonds (B,E) Permanent bonds (C,F) Sliding bonds. For the number fraction graphs, the sum of all of the generations is shown with a horizontal green dashed line. The deformation stretch is shown with a red dashed line on reference configuration graphs. Note that **Π**^*α*^ in general is a second-order tensor, and we used a one-dimensional representative for more convenient illustration. (For the references to color in this figure legend, the reader is referred to the web version of this article.)

The sliding only occurs when the sliding condition is met. Otherwise, bonds do not break and due to intrinsic hyperelasticity assumption they behave similarly to permanent bonds. The sliding condition (analogous to yield condition) is defined as

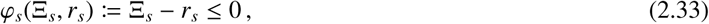

where Ξ_*s*_ is the sliding variable and *r_s_* is the sliding threshold, where all are scalar. The sliding variable is a function of deformation, or equivalently the invariants of deformation to guarantee its coordinate-invariance:

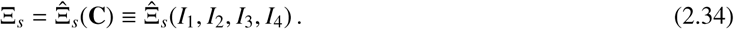

The sliding threshold *r_s_* is defined as the maximum value of Ξ_*s*_ obtained in the history of deformation (Simo, 1987)

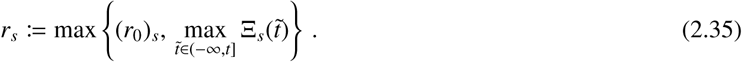

The initial value of *r_s_* before any loading is the initial sliding threshold denoted by 
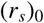
, which is a material property. Equations (2.33) and (2.35) indicate 
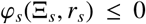
, which corresponds to the physically admissible deformations (Simo & Hughes, 1998). Sliding can only be initiated when *φ*_*s*_ = 0 (necessary condition). To determine the su cient condition, we need to include the loading direction by defining the second order tensorial normal to the sliding surface (*φ*_*s*_ = 0)

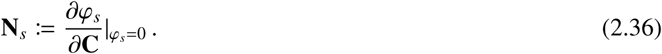

While the necessary condition for sliding is in place, two scenarios of loading can occur (Naghdi & Trapp, 1975)

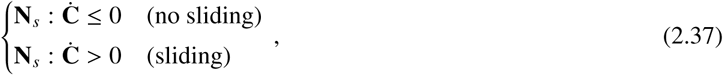

where only the second scenario can initiate a sliding behavior. Here, **Ċ** is the material time derivative of the right Cauchy-Green deformation tensor 
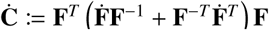
. When the necessary and sufficient conditions of sliding (*φ*_*s*_ = 0 and **N**_*s*_: **Ċ** > 0) are in place, the incremental sliding can be calculated using the sliding rule:

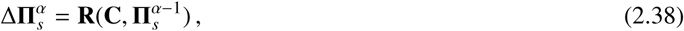

where **R** is a second-order tensor function. An example of sliding rule for a medium with a dominant fiber direction would be

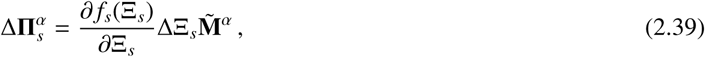

where *f_s_* is the sliding function, ΔΞ_*s*_ is the difference between sliding variables of two consecutive generations involved in the sliding process (Eq. (2.32)), and 
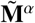
 is a second-order structure tensor. To define 
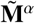
 it was assumed that the initial dominant fiber orientation is preserved in the sliding configurations, i.e. 
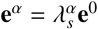
. Therefore, one can show (see Appendix B)

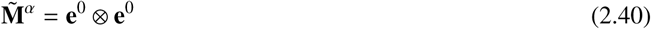

This particular constitutive relation indicates that the configuration of increments of sliding depends on the extent of deformation and the orientation of bonds in the master reference configuration. As another example, for three-dimensional cases where an analytical expression is available for **F**, one can choose 
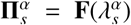
 (Rajagopal & Wineman, 1992). For completeness, because the reference configuration is time invariant after formation of bonds and the sliding process is incremental (Eq. (2.32)), the rate of sliding for each reforming bond generation is determined using

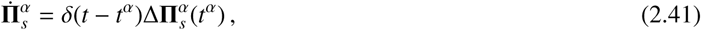

where *δ*(.) is the Dirac delta function that denotes the incremental nature of sliding process. This relation will be helpful later in the analysis of the second law of thermodynamics (Section 2.6).

### 2.5. Damage of bonds

Damage is an irreversible process that is due to a loss of chemical bonds (Krajcinovic, 2000). This is often observed as a softening behavior that is also referred to as Mullin’s effect (Mullins, 1969). The idea of relating damage to molecular bonds in tissue can be traced back as early as (Chu & Blatz, 1972) for modeling the hysteresis effect. In the current framework, damage of the various bonds (*D*) was considered as another ISV, where with a damaging load, the broken bonds may not be reformed, which in turn decreases the number of active bonds. That is,

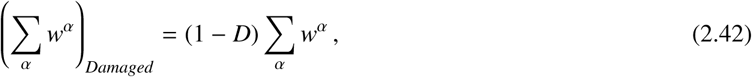

where *D* is the damage parameter of a certain bond type and it takes a value between zero and one (*D* ∈ [0, 1]) (Nims et al., 2016). Damage can be applied to each of the bond types (Fig. 4). By doing so, the total energy of the bonds decreases as the number fraction of damaged bonds increases. Hence, the free energy is

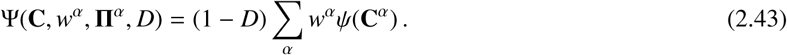

which always has a value less than the non-damaged case.

To determine the evolution of damage, similar to the sliding case, the damage condition is defined as

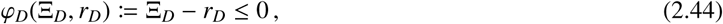

Here, 
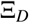
 is the damage variable that is a function of deformation invariants

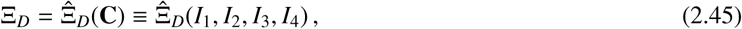

**Figure 4.**
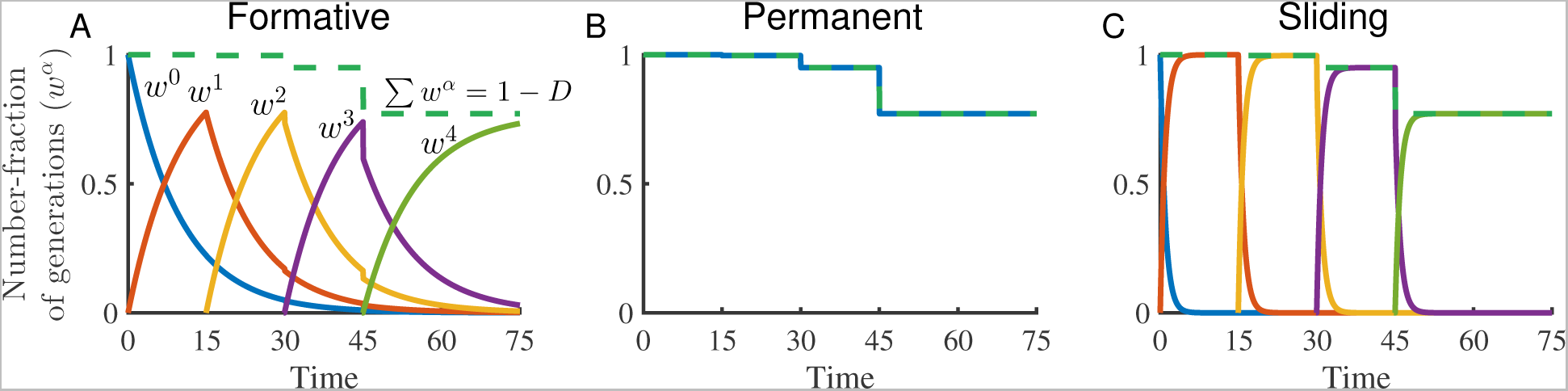
Effect of damage on kinetics: Multiple generations initiated at four consecutive step deformations at *t* = 0, 15, 30, 45 with a first-order rate equation with damage shown for (A) formative bonds (B) permanent bonds and (C) sliding bonds, the sum of bonds declines after each step of loading until the final value of 0.75 (*D* = 0.25). (For the references to color in this figure legend, the reader is referred to the web version of this article.)

and *r_D_* is the damage threshold defined as the maximum value of Ξ_*D*_ in the past history of deformation (Simo, 1987). The damage threshold *r_D_* is defined separately for each bond type as

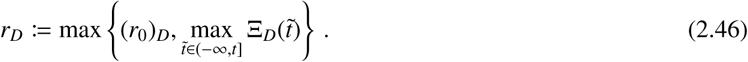

The value of *r_D_* before any loading is the initial damage threshold 
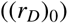
, which is a material property. Damage occurs when *φ*_*D*_ = 0 (necessary condition). The loading direction determines the su cient conditions. By defining the second order tensorial normal to the damage surface (*φ*_*D*_ = 0) as

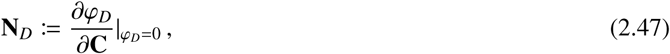

and by using the following normality conditions (Simo, 1987)

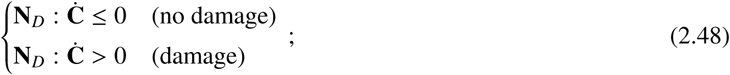

damage only occurs when (*φ*_*D*_ = 0 and **N**_*D*_: Ċ > 0). Thus, during a damaging load, the damage parameter increases according to the damage rule

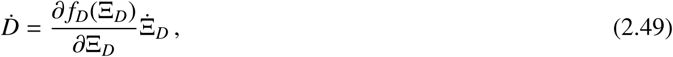

where *f_D_* (Ξ_*D*_) is the damage function. Note that the damage variable Ξ_*D*_ is different from damage parameter (*D*), where *D* is the number fraction of permanently broken bonds.

### 2.6. Implications of the second law of thermodynamics

Any physical process must comply with the second law of thermodynamics, thus we check for compatibility of reactive inelasticity with the second law of thermodynamics in terms of the Clausius-Duhem inequality. By ignoring the thermal effects, the local form of Clausius-Duhem inequality reads as (Coleman & Gurtin, 1967)

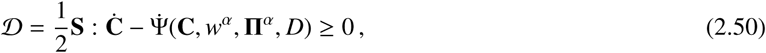

where 
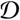
 is the instantaneous local dissipation of energy, and for convenience will be referred to as dissipation. Here, (:) is the double contraction ((**A:B**)_*i j*_ = *A_i j_ B_i j_*) and 
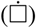
 stands for material time derivative (Holzapfel, 2000). Using the chain rule of differentiation and Eq. (2.11) we get

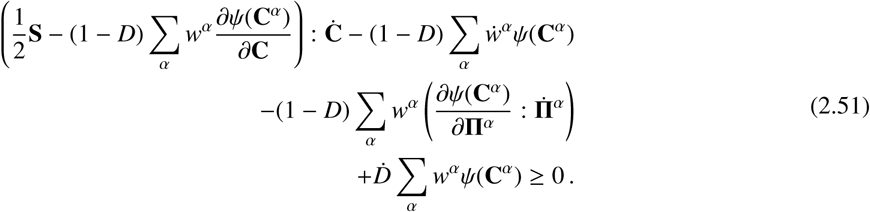

Thus, to have a physically admissible process in the most general case each of the following conditions should be separately satisfied (Coleman & Gurtin, 1967; Murakami, 2012)

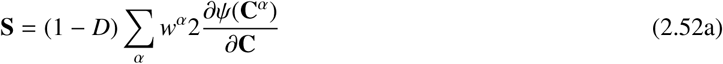

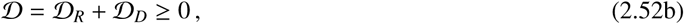

where

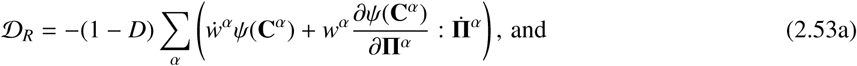

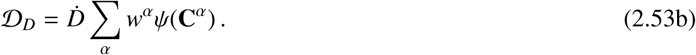

The first condition (Eq. (2.52a)) results in a new form of Eq. (2.12) where stress is scaled with damage. The second condition (Eq. (2.52b)) describes the dissipation of energy in a reactive inelastic process, where 
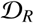
 is dissipation due to reactive bond breaking/reformations and change in reference configuration, and 
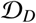
 is the dissipation due to damage. The detailed proofs of the following results are included in Appendix C, and the following are the final results of reactive dissipation for each bond type.

We consider first only the dissipation due to bond breaking and reforming. For formative bonds, dissipation is always positive because the breaking bonds (*α* < *N*) have a negative breakage rate, and the reforming bonds (*α* = *N*) are energy-free. That is,

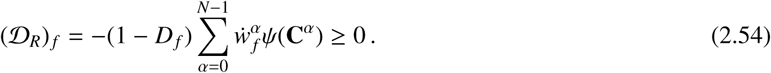

The permanent bonds dissipate no energy because there is no breakage and reformation occurring for these bonds:

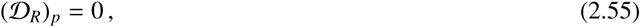

and for sliding bonds dissipation is

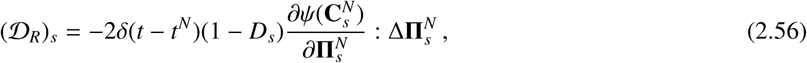

where the Delta function form occurs due to the rate of sliding that was shown in Eq. (2.41). Half of this dissipation term (Eq. (2.56)) is due to the process of breakage and reformation, and the other half is due to a shift in the reference configuration. Accordingly, when the following condition is in place, the Clausius-Duhem inequality is satisfied

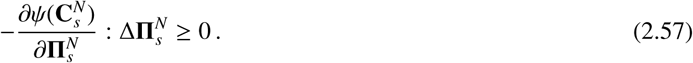

Equation (2.57) implies that at a given **C**, an increment of sliding only serves to reduce the energy level or equivalently to dissipate energy compared to a non-sliding case. Another implication of this condition is that once sliding has dissipated all of the energy in the system, no further plastic deformation can occur and no further energy can be stored in the bonds.

Considering the case where damage is involved 
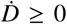
 and since the energy and number fraction of bonds have positive values, the second term of Eq. (2.52b) is always positive. Therefore,

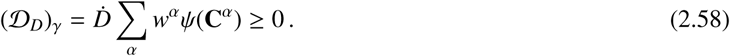

Note that the above relation is only valid when there is no process of biological recovery and growth that may be present in living systems, where special considerations should be employed that are beyond the scope of this study.

In conclusion, as a consequence of the second law, formative and permanent bonds are ‘second-law compatible’ for a valid kinetics relation, and for sliding bonds, the change in reference configuration (Eq. (2.39)) needs to be defined so as to satisfy Eq. (2.57). For damage, *f_D_* (Eq. (2.49)) needs to be selected to guarantee 
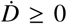
 (e.g., cumulative distribution function of any continuous statistical distribution). Examples of appropriate constitutive relations will be provided in Section 3.

### 2.7. Summary of formulation

In summary, a reactive inelasticity model was formulated that defines different types of bonds corresponding to different mechanical behaviors. The combination of bond types should be selected according to the desired mechanical behaviors, where formative bonds should be used for a transient behavior with zero equilibrium stress, permanent bonds for a hyperelastic response, and sliding bonds should be used when a plastic deformation is occurring. Damage may be added to each of the bond types, where it reduces the number fraction of active bonds, and thus reduces the ability of material to absorb energy. In selecting the constitutive relations it is perhaps more convenient to take the discrete steps that are outlined in Table 2.1. In the following section, several illustrative examples are provided that both help in understanding the model and also to exemplify some practical applications to mechanics of soft tissue, where reactive inelasticity can be useful.

**Table 2.1.** Summary of structure of a reactive bond type’s constitutive relations.

|  |  |
| --- | --- |
| Step 1 | Select the bond types (i.e., formative, permanent, sliding) |
| Step 2 | Select the kinetics relation (e.g., Eq. (2.27)) |
| Step 3 | Select the intrinsic hyperelasticity relation $\psi_\gamma(\mathbf{C}^\alpha)$ (e.g., neo-Hookean) |
| Step 4 | If sliding bonds, select appropriate constitutive relations:<br>Sliding variable ($\Xi_s$) (Eq. (2.34))<br>Sliding rule and function ($f_s$) (Eq. (2.39)) |
| Step 5 | If damage is included, select appropriate constitutive relations:<br>Damage variable ($\Xi_D$) (Eq. (2.45))<br>Damage rule and function ($f_D$) (Eq. (2.49)) |

## 3. Illustrative examples

To demonstrate key features of the reactive inelasticity framework we provide four numerical examples. First, the sensitivity of formative bonds to the kinetics rate is demonstrated using a Heaviside step loading that shows transitions between different bond types (Example I). Second, we compare the stress response of the bond types to the classical models; in particular the response of formative bonds is compared to quasi-linear viscoelasticity (QLV), permanent bonds to theoretical neo-Hookean hyperelastic material, and sliding bonds to a deformation-based plasticity model (Example II). Third, we simulate the stress response of an increasing cyclic loading by using a fiber-exponential constitutive relation that is commonly used in tissue mechanics, and we compare the stress response between non-damaged and damaged cases (Example III). Finally, incremental stress relaxation with softening is presented, which has importance to understand mechanisms of plasticity and damage in tissue. This is done by using a combination of formative bonds and either sliding bonds or permanent bonds with damage (Example IV). The model is implemented as a Matlab function using a custom written code intended for uniaxial deformations. The source code and details of implementation are provided as supplementary material and are accessible via the following online repository (Safa, 2018).

For all of the examples, the axial component of the Cauchy’s stress tensor (i.e., *T* = *T*_33_) was used to represent the state of stress during a uni-axial isochoric deformation

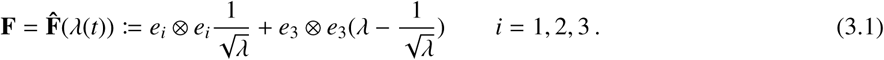

The following constitutive relations were implemented in the steps outlined in Table 2.1:

- **Step 1- Bond type selection:** The bond type was selected depending on the expected mechanical behavior. Formative bonds were used for a viscoelastic behavior with zero equilibrium stress, permanent bonds for a hyperelastic behavior, and sliding bonds were used for plastic deformation. Note that for Example IV, which includes a combination of bonds, the response was calculated independently for each bond type and added according to Eq. (2.11).
- **Step 2- Kinetics relation:** For all of the examples, a first-order kinetics relation was used (Eq. (2.27)).
- **Step 3- Intrinsic hyperelasticity:** Three types of non-linear constitutive relations were used. First, we used the neo-Hookean relation in Examples I and II. This is the simplest non-linear constitutive relation and was used to demonstrate the general behaviors of the model, where the potential energy is

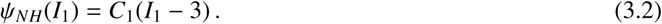

Here, *C*_1_ is the independent model parameter with dimensions of stress.

Second, for relevance to tissue applications, in Examples III and IV we used a one-dimensional exponential constitutive relation with a non-zero sti ness only in the *e*_3_ direction (fiber direction) in tension (Jacobs et al., 2014; Schmidt et al., 2014)

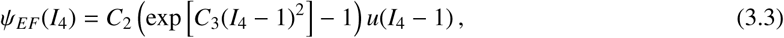

where *C*_2_ and *C*_3_ are positive-valued model parameters.

Third, for showing a three-dimensional tissue mechanics application, we used a Holmes-Mow material (Example IV)

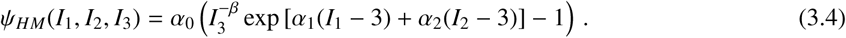

In this relation, *α*_0_ is a positive number with the dimension of stress and [*β*, *α*_1_, *α*_2_] are positive non-dimensional numbers. To comply with the energy‐ and stress-free reference configuration, *β* = *α*_1_ + 2*α*_2_ must be satisfied (Holmes & Mow, 1990). It also should be noted that the Holmes-Mow model parameters are related to the more familiar infinitesimal linear-elastic parameters as:

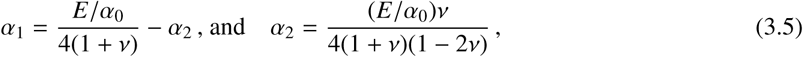

where *E* is the Young’s modulus, and *v* is the Poisson’s ratio. Hence, [*E, v*] were used in the following instead of the original Holmes-Mow parameters.

- **Step 4- Sliding:** The overall stretch was selected as the sliding variable

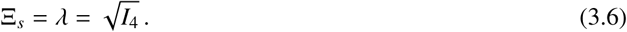

To describe the sliding rule for the one-dimensional fiber intrinsic hyperelasticity relation (Example II) (Eq. (3.3)) the form represented in (Eq. (2.39)) was employed with the sliding function as

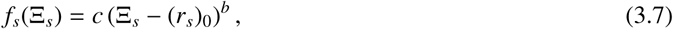

where the constant parameters [*c, b*] are dimensionless positive numbers, and 
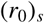
 is the initial sliding threshold. Also, e_3_ was taken as the sliding direction and was used to define structural direction tensor as 
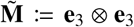
 (Eq. (2.39)). In this case, as long as 
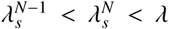
the condition of Eq. (2.57) is satisfied, and it can be easily verified that 
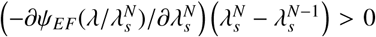
, which proves the compatibility of these constitutive relations with the second law. For the three-dimensional constitutive relation cases (Examples I,II,IV) it was assumed that 
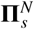
 takes the form of 
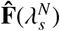
 during sliding, where by using the Taylor series expansion, it can be shown that all of the constitutive equations are compatible with the second law of thermodynamics (for details see Appendix C).

- **Step 5- Damage:** For the case with damage of bonds, similar to sliding, the overall stretch was used as the damage variable

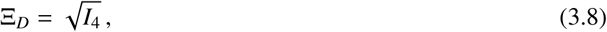

and a Weibull’s cumulative distribution function was used as the damage function (Nims et al., 2016)

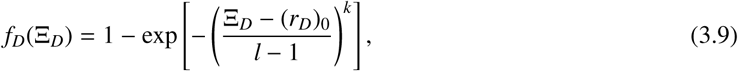

where *k* is the shape parameter, *l* is the scale parameter, and (*r*_0_)_*D*_ is the initial damage threshold.

### 3.1. Sensitivity of the formative bonds to the kinetics rate (Example I)

As discussed in Section 2, the kinetics rate is the predominant difference between various bond types. To show this, we simulated the response of formative bonds to a Heaviside step deformation at a wide range of reaction kinetics rate, where the only variable was the time constant (*τ*_*f*_) (Fig. 5). For a large *τ*_*f*_ (slow kinetics rate) the response approaches the hyperelastic behavior (permanent bonds), and as *τ*_*f*_ gets smaller (high kinetics rate) the response approaches a singular stress behavior (sliding bonds). For all of the kinetics rates, the response at *t* = 0 is the same and equal to the hyperelastic stress for that deformation. This response at *t* = 0 also corresponds to the peak stress response of the initial generation in Eq. (2.24) and (2.25).

This simple example lies at the heart of reactive inelasticity and demonstrates the consistency of inelastic behaviors that are all based on the kinetics of breaking and reforming bonds. In what follows, more complex loading scenarios are investigated that are all based on a summation of incremental step deformations.

### 3.2. Response of various bond types and comparison to classic models (Example II)

Example II is the stress response of the three bond types to a simple loading/un-loading deformation (Fig. 6A) with a neo-Hookean constitutive relation for intrinsic hyperelasticity. Formative bonds show stress relaxation, which is a common characteristic of a viscoelastic response (Fig. 6B, E). Permanent bonds show a one-to-one relation between deformation and stress with no history dependence (hyperelastic) (Fig. 6C, F). Sliding bonds show a difference between loading and unloading, where during loading, the state of zero-stress was shifted, which indicates a plastic deformation (Fig. 6D, G).

We compared the response of the three bond types to classic models (model parameters are detailed in Table D.1). In particular, the response of formative bonds is compared to quasi-linear viscoelasticity (QLV) (Fig. 6B,E) (Fung, 1993), where

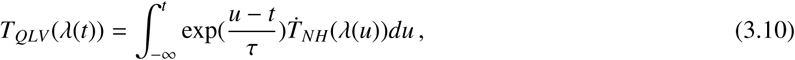

and 
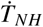
 is determined using Eq. (2.20) and (3.4). The response of permanent bonds is compared to the theoretical hyperelastic response of a neo-Hookean material (Fig. 6C, F). Lastly, the response of sliding bonds is compared to a simple deformation-driven plasticity model (Fig. 6D, G), where

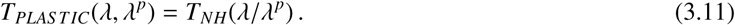

Here *λ* corresponds to stretch, also note that the superscript *p* indicates the plastic part of deformation. For post-yield loading *λ*^*p*^ is calculated as

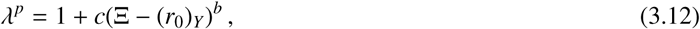

where Ξ = Ξ_*s*_ and 
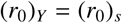
, also [*c, b*] have the same values as in the sliding constitutive relations.

**Figure 5.**
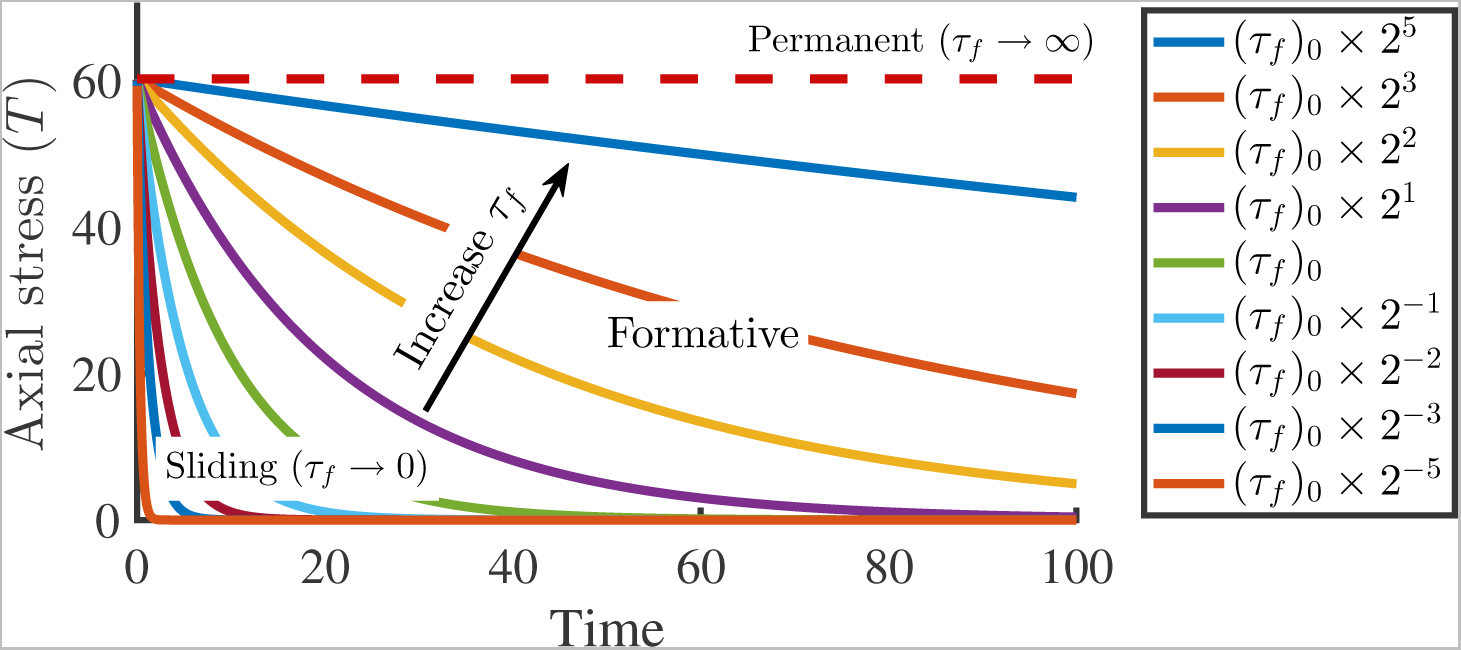
Example I: Time response of formative bonds to a Heaviside-step deformation in a large range of kinetics. The kinetics rate is controlled by time constant of reaction where (*τ*_*f*_)_0_ = 10 and the range of time constants is covered by multiplying (*τ*_*f*_)_0_ with powers of two. Formative bonds show an asymptotic decay in Cauchy stress during a sustained loading, whereas for large values of *τ*_*f*_ (slow kinetics rate) the response approaches a hyperelastic behavior (permanent bonds) marked with dashed red line, and at small *τ*_*f*_ (high kinetics rate), the stress response approaches a singular behavior that is immediately decays to zero (sliding bonds). The step deformation is *λ*(*t*) = 1:1*u(t)* and the neo-Hookean intrinsic hyperelasticity parameter is *C*_1_ = 100. (For references to color in this figure legend, the reader is referred to the web version of this article.)

Reactive inelasticity successfully reproduced each of the classical models. Note that for large deformations, the conclusions are valid for permanent bonds; however, the behavior of formative bonds and QLV slightly deviate, which is due to the Boltzmann’s superposition principle in QLV (Ateshian, 2015). For plastic deformation, although there is no difference between the stress response of deformation-based plasticity model and reactive sliding, the two approaches indicate different physical meanings, where by simply shifting the reference configuration (as in the deformation-based formulation) the process of breakage and reformation is not considered. Mechanistically, an inelastic bond sliding should occur as an explicit consequence of a bond dissociation and reattachment, and thus we suggest that the reactive interpretation of this process is physically appropriate. From these examples, it is evident that the reactive inelasticity framework is capable of reproducing various mechanical behaviors, where the same sets of constitutive relations are used for each of the bond types.

### 3.3. Cyclic loading and damage (Example III)

Example III demonstrates the response of single bond types with the model parameters listed in Table D.2 to a cyclic loading (Fig. 7A). In response, formative bonds dissipate energy in a cyclic hysteresis (Fig. 7B), while the permanent bonds have a hyperelastic response that is independent of history of deformation (Fig. 7C). The sliding bonds undergo plastic deformation that is evident by the shift in the point of zero stress after each cycle, while there is no change in sti ness during the consecutive loading parts (Fig. 7D). Note that the sliding bonds account for strain softening that results in a decrease in stress after reaching the peak stress (points 3 and 4 in Fig. 7D), and it is different from strain hardening that is associated with plasticity in metals and some engineering polymers. Further, by adding damage (decreasing the total number of bonds) the Mullin’s effect was captured for all three types of bonds. In addition to decreasing sti ness and the corresponding stress values, the damage increases the difference between the loading and unloading parts of the stress response. This results in increased hysteresis and further dissipation of energy (Fig. 7E, F, and G).

The expansion of sliding and damage thresholds are shown in Fig. 7A by the dashed lines. It is evident from the simulations that the behavior of bonds, especially permanent bonds with damage and sliding bonds are similar. However, certain characteristics are unique to each bond type. For example, contrary to sliding bonds, there is no shift in reference configuration for permanent bonds with damage. Importantly, from an experimental perspective, it is not possible to distinguish the mechanical response of plastic deformation from that of damage in loading, and their difference becomes evident only in unloading.

**Figure 6.**
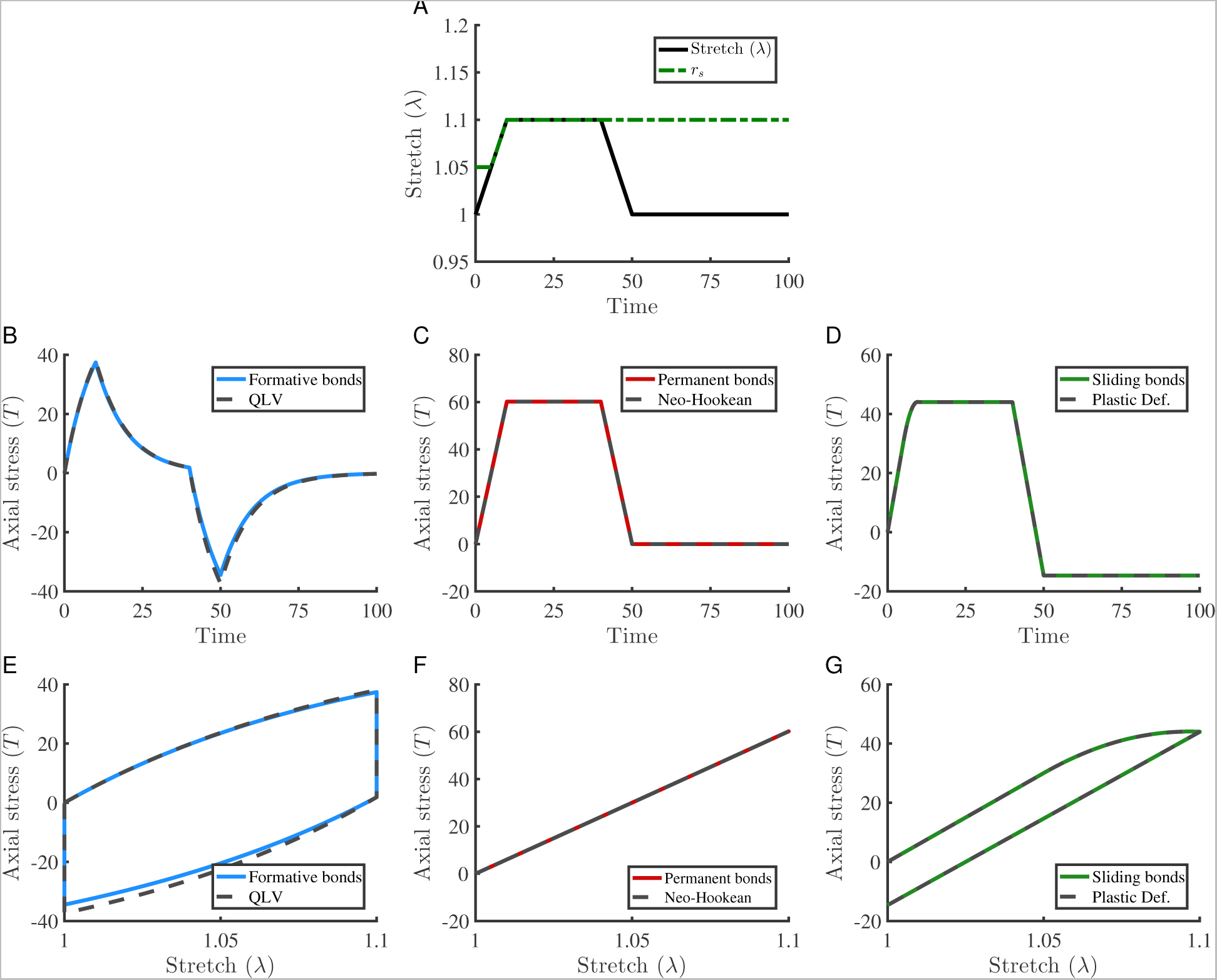
Example II: Mechanical response of different bond types to a simple loading/un-loading deformation. (A) Stretch profile of loading, and evolution of the sliding threshold. In (B,E) formative bonds were compared to quasi-linear viscoelasticity, (C,F) compares permanent bonds to the theoretical stress response of a neo-Hookean material, and (D,G) shows the comparison between sliding bonds’ response and a plasticity model in deformation space.

### 3.4. Incremental stress relaxation with softening (Example IV)

In Example IV a typical response of tissue in a stress relaxation test is demonstrated that also includes softening. Softening in tissues often occurs at high deformations (Szczesny & Elliott, 2014; Lynch et al., 2003). It is not possible to distinguish between damage and plastic deformation solely by loading the tissue in tension and analyzing the stress response. This is demonstrated in the example by using a combination of formative bonds and either sliding bonds, or permanent bonds with damage (Fig. 8). In these examples, formative bonds account for the transient stress response, and sliding (or permanent bonds) produce the equilibrium part of the stress response (model parameters in Table D.3). Identical behaviors are observed during ramp and relaxation loading phases (Fig. 8). However, during unloading it is evident that the reference configuration is shifted when implementing the sliding bonds, where for permanent bonds with damage there is no shift in the zero-stress configuration.

Experimental evaluation of this shift in the reference configuration in soft tissue is often challenging due to nonlinearity in stress-strain behavior, when the initial sti ness is low that gradually increases by further deformation (toe-region effect). However, by having a theoretical framework for both of the candidate mechanical behaviors (i.e., plastic deformation and damage) it is possible to fit the models to the loading phase and make predictions about either the unloading phase, or other experimental results (e.g., cyclic loading in Example III) to assess the success of the model in predicting the experimental data, and thus evaluating the contribution of each of the aforementioned inelastic behaviors in tissue.

**Figure 7.**
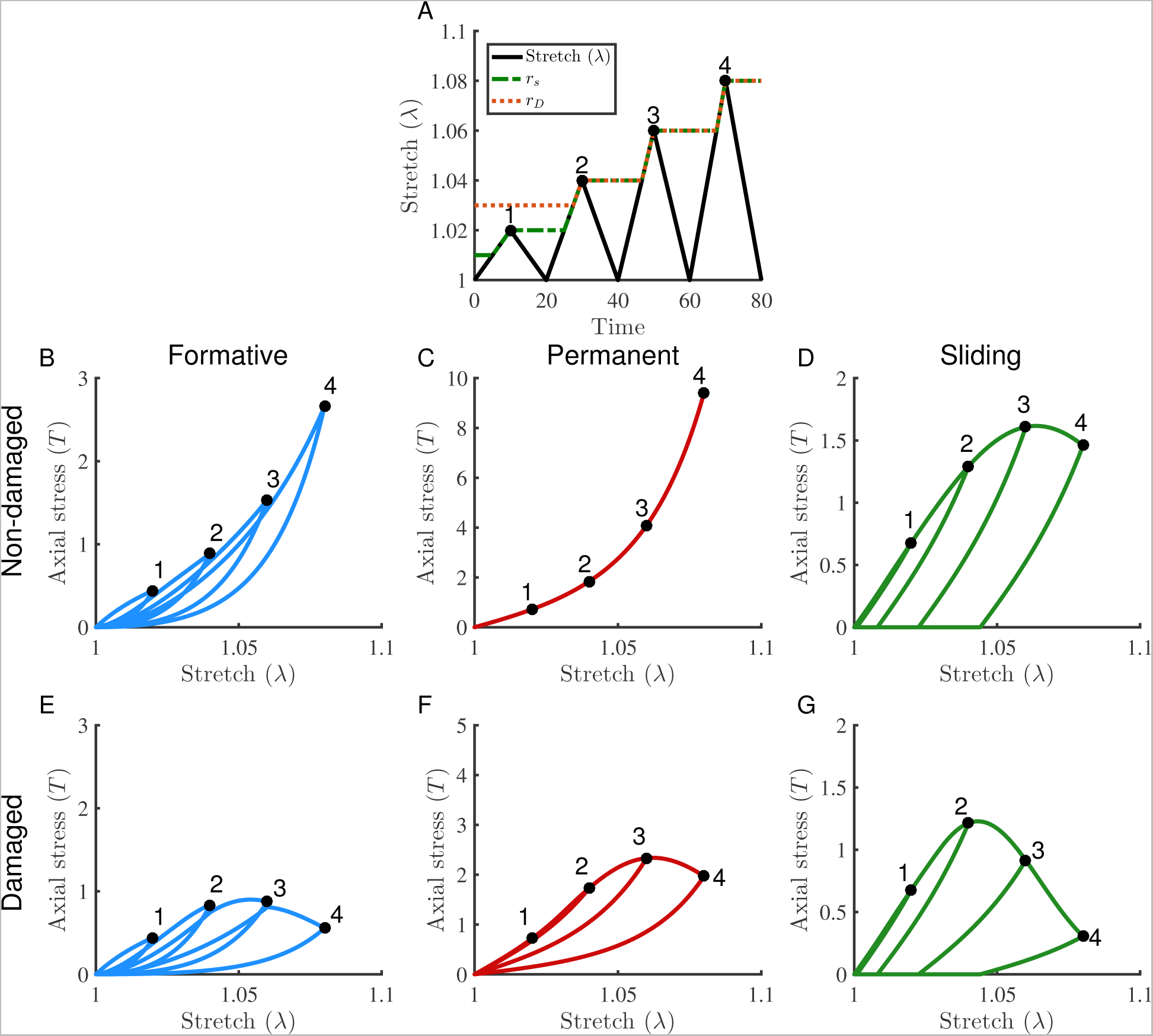
Example III: Cyclic loading and damage. (A) an increasing cyclic loading protocol applied for non-damaged (B) formative, (C) permanent, (D) sliding bonds. When damage is added to the response for (E) formative, (F) permanent, and (G) sliding bonds, all the bond types show a softening behavior. The evolution of sliding and damage thresholds is also shown in (A) in response to loading and unloading phases, where sliding starts after 
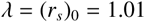
, and damage after 
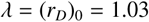
.

## 4. Discussion and concluding remarks

This study provides a structurally inspired framework for modeling inelasticity in tissue. It uses a thermodynamic scheme, where internal state variables and kinetics of molecular bonds are employed to capture the major inelastic behaviors of tissue. This framework consistently addresses a range of inelastic behaviors. The formative, permanent, and sliding bonds correspond to viscoelastic behavior with zero equilibrium stress, hyperelastic response, and plastic deformation, respectively. Additionally, the formulation of bond kinetics allows for the inclusion of damage by reducing the number of bonds, which introduces damage as an inelastic behavior to all of the bond types. Any combination of the bond types can be employed to model various kinds of inelastic behaviors, such as a combination of formative bonds and permanent bonds with damage to model a viscoelastic behavior with softening.

**Figure 8.**
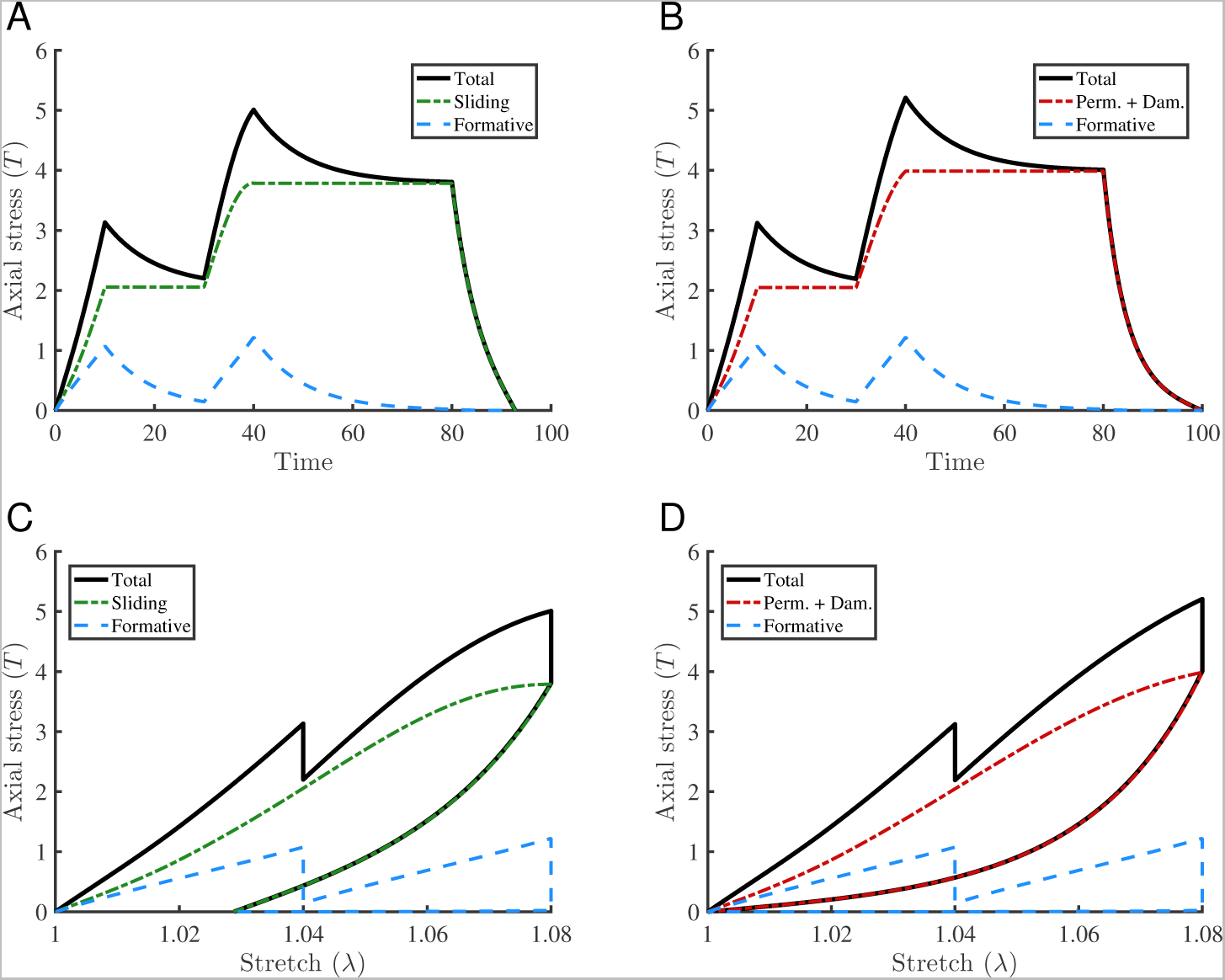
Example IV: Incremental stress relaxation with softening for (A,C) formative and sliding bonds, and (B,D) formative bonds with permanent bonds and damage. The effect of selection of either sliding bonds or permanent bonds is only observed during unloading. When the sliding bonds are used there is a shift in the reference configuration (C) which is not true for permanent bonds with damage (D).

The reactive inelasticity framework is structurally inspired, thus the concept of bonds and their energy level have physical relevance—unlike the majority of the phenomenological models. Connective tissue’s matrix is mostly made of cross-linked collagen. These molecules are cross-linked to each other via several mechanisms including enzymatic lysyl oxidase-mediated and non-enzymatic glycation-induced cross-links to produce larger collagenous structures (Reiser et al., 1992; Eyre et al., 2008). The mechanical properties of collagenous material are sensitive to the density, structure and properties of these cross-links (Depalle et al., 2015). When overloaded, collagen fibrils show local deformities (e.g., kinks) that contain denatured-collagen-rich regions (Herod et al., 2016; Veres et al., 2014). Recent advancements in using collagen hybridizing peptides (CHP) provides opportunities to visualize the spatial distribution and level of disruption in these regions after mechanical overloading (Li & Yu, 2013; Zitnay et al., 2017). These observations suggest that there is a strong correlation between mechanical loading and the state of molecular bonding. Two potential mechanisms of molecular disruption, shear and tension dominant, were suggested based on molecular dynamics experiments that closely resemble the sliding and damage processes (Zitnay et al., 2017). Despite these advancements in visualizing the molecular state of tissue and its relation to their mechanical behavior, further experimental work needs to be done to elucidate structural mechanisms of inelasticity in terms of molecular bonding.

The reactive inelasticity framework is a generalization of the reactive viscoelasticity model (Ateshian, 2015), where an inclusive and consistent treatment of inelastic processes is provided. The reactive viscoelasticity model of Ateshian provided the essential concepts of weak and strong bonds that correspond to formative and permanent bonds, respectively. We chose the terminology of formative and permanent bonds because of their analogy to kinetics rate of bonds and their behavior. Additionally, in contrast to (Ateshian, 2015) we based our formulation on the number fraction of bonds rather than mass fraction, which emphasizes that in general, bonds are levels of energy and individual bonds do not correspond to a physical mass. Although at the first sight these parameters appear to be equivalent, due to their non-dimensional nature, this non-inertial bond assumption is particularly important when considering the singular accelerations arising in the case of sliding bonds. Massless bonds keep the model in the realm of admissibility in the general sense. Further, by making these generalizations we showed that the bonds are interchangeable at the extremes of kinetics rate, where the behavior of formative bonds is hyperelastic (permanent bonds) in the limit of slow kinetics rate (Fig. 5), and at high rates of bond kinetics formative bonds turn into sliding bonds that correspond to a plastic deformation (Figs. 6 and 7).

Plastic deformation has been experimentally observed in tissue at the macro‐(Maher et al., 2012), micro‐(Szczesny & Elliott, 2014; Lee et al., 2017), and nano-scale (Shen et al., 2008). Several phenomenological ISV modeling approaches have been taken to address these phenomena that often are focused to address specific inelastic behaviors and usually omit the relationships between different inelastic behaviors and their mechanisms (Zhang & Sacks, 2017; Fereidoonnezhad et al., 2016; Peña, 2014; Maher et al., 2012). In the current framework, we particularly focused on providing a consistent formulation between different inelastic behaviors and their structural relevance. In a similar manner, for cross-linked polymers a molecular bond approach has been used to model stress relaxation and plastic necking instability (Meng & Terentjev, 2016; Meng et al., 2016); however, in that theory, only plastic necking rather than general plastic deformation was addressed, and therefore, their formulation does not explicitly provide for the shift in unloaded configuration that is observed in tissue, such as non-recoverable interfibrillar sliding (Lee et al., 2017). In our model, the generation reference deformation gradient (**Π**^*α*^) is a state variable that is directly related to the plastic deformation and thus is an observable variable accounting for the shift in unloaded configuration of the material.

Damage was applied to each of the bond types regardless of their kinetics rate. This allowed for modeling the effect of damage on different components of structure that produce the mechanical behavior of material, such as degradation in the viscous components of a viscoelastic behavior without a ecting its elastic part. Previously, a similar concept was used to study damage in cartilage that was only applied to the permanent bonds. This resulted in the damage effect being limited to the equilibrium behavior (Nims et al., 2016). The reduction in number fraction of active bonds results in a decrease in the ability of material to absorb energy, which provides a generalized definition of damage as a behavior. By adding this ability, it is possible to model mechanical behavior up to rupture, where, if the formative bonds could not be damaged, no rupture could occur. Another benefit of this definition is in that it forms a clear theoretical basis for distinction between plastic deformation and damage. As shown in Fig. 7, plastic deformation alone does not result in a decrease in sti ness; however, adding damage would cause a reduction in sti ness, which is a common mechanical property used to assess the extent of damage in a material. By using the current theoretical framework for addressing damage and other forms of inelastic response that was provided in this study, it is possible to elucidate the mechanisms and role of each inelastic behavior in the mechanical response of tissue from the experimental stress response of the material. One future direction, for example, is using a specific loading profile with repeated scenarios (e.g., loading, unloading and reloading) to curve fit different versions of reactive inelasticity and using the fit parameters to predict an independent experimental mechanical behavior. From the quality of the predictions, one can assess the relevance of the involved inelastic behaviors and their contribution.

In this framework, one of the limitations may be that we made a continuum body assumption to relate the molecular energy to the macro-scale mechanical behavior, where hierarchical structural effects, as well as statistical considerations for molecular bonding were not included; however, the simplicity of the model and its ability to reproduce a spectrum of macro-scale behaviors were chosen here over complexity of formulation. Additionally, we modeled the plastic deformation by using a finite-strain framework with a constrained direction, and damage was modeled as a scalar variable that may overly constrain a generalized three-dimensional case (Ju, 1990). These issues can be addressed by adding multiple bond types with different directional preferences to achieve a desired spatial characteristic (Nims et al., 2016). Finally, since tissues have a significant fluid content, the fluid pressurization and flow can play an important role in the mechanical response. Those effects were not included in this model, which is a limiting factor for applying this framework to experimental data. However, the effect of fluid content can be added as in a multiphasic mixture model with inelastic solid phase that uses reactive inelasticity (Huang et al., 2001).

In conclusion, we have provided a structurally inspired framework for modeling inelastic behaviors of tissue based on the kinetics of reactive molecular bonds that break and reform in response to mechanical loading. The model addresses viscoelasticity, plastic deformation, and damage in an inclusive framework. We introduced three types of molecular bonds: formative, permanent, and sliding that result in viscoelastic, hyperelastic, and plastic deformation behaviors. Damage was applied to each of the bond types by reducing the number fraction of bonds, which consequently decreases the ability of material to absorb energy. All of the aforementioned inelastic behaviors are modeled within the same framework and by using similar sets of constitutive equations. This allows for investigating the mechanisms of inelasticity by providing a comprehensive theoretical basis for analyzing the mechanical response of tissue that can be used to understand mechanical loading induced pathological conditions, to develop more effective treatment protocols, and to engineer replacement tissues.

## 6. Acknowledgment

Research reported in this publication was supported by the National Institute of Biomedical Imaging and Bioengineering of the National Institutes of Health under award number R01EB002425. The content is solely the responsibility of the authors and does not necessarily represent the official views of the National Institutes of Health.

## Appendix A. Stress and relative deformation

By using the definition of free energy of the system from Eq. (2.11) the second Piola-Kirchho stress tensor for a bond type in terms of number fraction of bonds (*w*^*α*^) and relative deformation (represented by relative right Cauchy-Green deformation **C**^*α*^) is

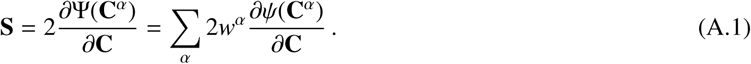

Using the chain rule of differentiation and Einstein’s index notation:

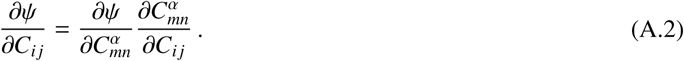

Using Eq. (2.2) and noting that the reference deformation gradient tensor (Π^*α*^) is fixed for a specific generation;

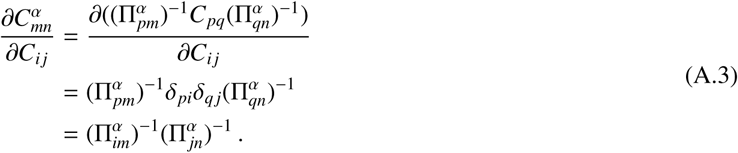

By substituting this result in Eq. (A.2) we get

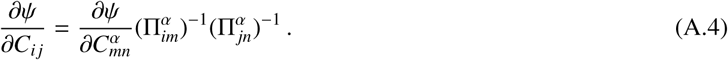

Or, in tensor notation

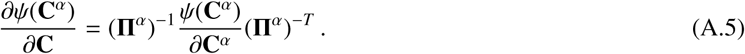

Substituting this result back in Eq. (A.1) gives

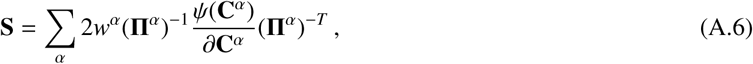

which provides the full form of the second Piola-Kirchhostress for a bond type in a reactive material, Eq. (2.13).

## Appendix B. Proof of the structure tensor 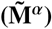 expression in Eq. (2.40)

By assuming that the stretch in the fiber direction for the reference configuration of the *α* generation is 
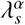
 which has a value that is infinitesimally different from that of generation (*α* - 1), and by using Taylor’s approximation one can show

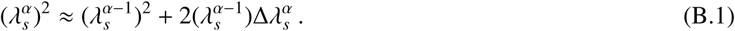

Additionally, based on Eq. (2.32) from the corresponding sliding configuration deformation gradients we have:

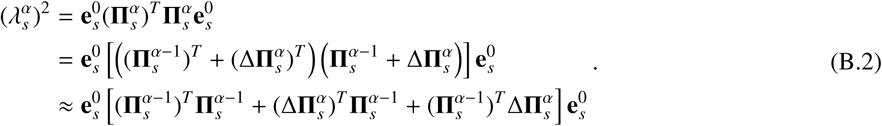

By combining Eqs. (B.1) and (B.2) and by substituting 
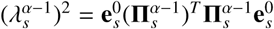
we get

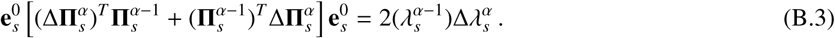

Therefore, by assuming 
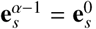
 one can easily show that

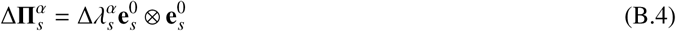

holds. Finally, by taking that 
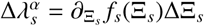
the expression 
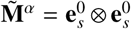
 is resulted in terms of Eq. (2.39). This relation indicates that during sliding, the dominant fiber direction is preserved and only the reference length changes. This relation can be enhanced by including the effect of permanent changes to the dominant fiber direction as seen in some fibrous tissues, such as the intervertebral disc (Dittmar et al., 2016), by using phenomenological relations to correlate 
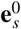
 and 
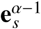
.

## Appendix C. Reactive dissipation due to reactive bond breaking and reformation

In the following, we investigate the compatibility of different bond types with the second law of thermodynamics. At a given time, consider a set of bonds from the same type that are grouped into breaking generations 
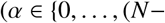

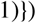

and a reforming generation (*α*= *N*), hence Eq. (2.53a) can be rearranged into

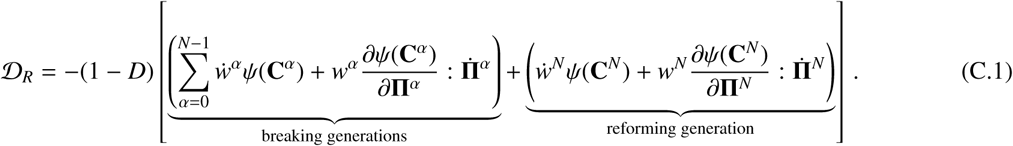

### Appendix C.1. Formative bonds

According to Eq. (2.27) for all of the breaking bonds we have

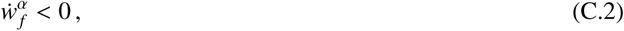

and we know in general that the free energy in a deformed state is considered positive

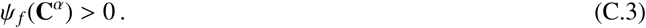

Additionally, the reforming bonds reform in an energy free state (Section 2.4.1)

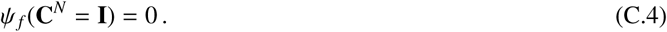

For the breaking formative bonds the reference deformation gradient is fixed 
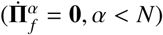
and for their reforming generation (*α*= *N*) (Eq. (2.30))

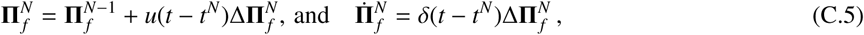

where *u(t)* denotes the unit step function, and *δ*(*t*) is the Dirac delta function. Further, using Eq. (2.28) one can show

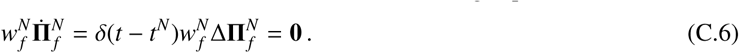

Therefore, for formative bonds

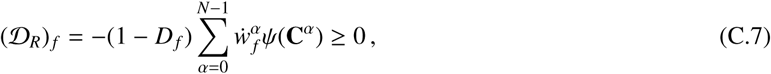

which indicates that during breaking and reformation of formative bonds, dissipation is positive and is due solely to breakage of bonds.

### Appendix C.2. Permanent bonds

Since for these bonds there is just one generation and there is no breakage and reformation, we have 
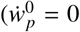
 and 
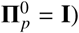
 as a result

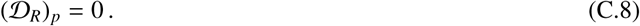

This result denotes that there is no dissipation of energy for permanent bonds due to a lack of reactive breaking and reformation.

### Appendix C.3. Sliding bonds

For these bonds, the only generations with non-zero dissipation terms are the last breaking (*α* = *N* - 1) and the reforming generations (*α* = *N*). Their number fractions and kinetics rates are (Section 2.4.3)

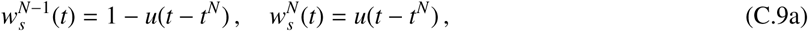

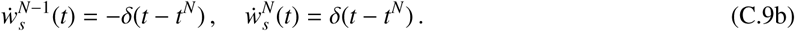

In addition, according to Eqs. (2.32) and (2.41) the corresponding 
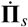
 for those bonds during sliding reads as

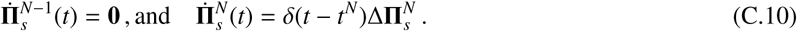

As a result

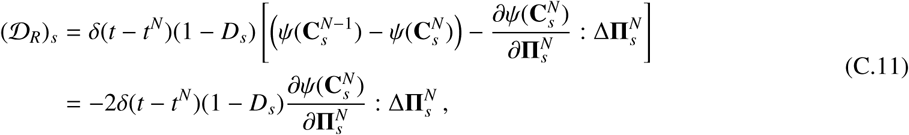

where we used the Taylor series to write 
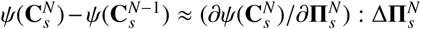
. Therefore, half of this dissipation is driven by the process of breakage and reformation, and the other half is explicitly due to a shift in the reference configuration. The above relation corresponds to a positive value for dissipation when

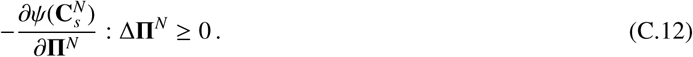

For the constitutive relations and deformation used in this paper, we can analytically verify the above inequality as follows.

#### Appendix C.3.1. Exponential fibers

Since the only non-zero component of the deformation and stress is in the fiber direction, a one dimensional formulation can be used for convenience, where 
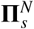
 is represented by 
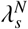
 As a result, the inequality (C.12) reads as

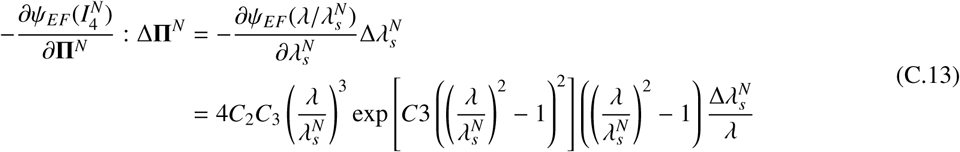

where 
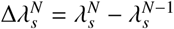
. As a result, since *λ^N−1^ < λ^N^ <λ* the above expression has a positive value.

#### Appendix C.3.2. Neo-Hookean

Because of the assumption that 
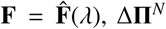
 can be written as 
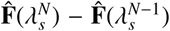
 and by using the Taylor’s expansion

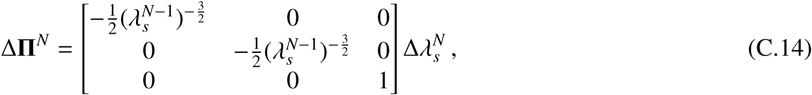

also, one can write

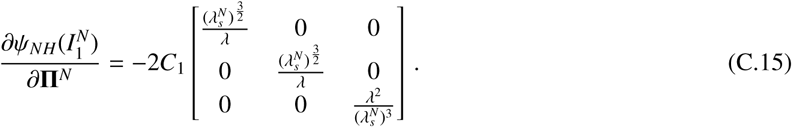

Therefore, the inequality (C.12) up to the first order relative to 
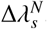
, reads as

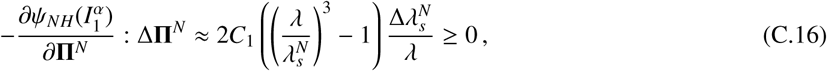

which is clearly valid when *λ^N−1^ < λ^N^ <λ*.

#### Appendix C.3.3. Holmes-Mow

The ΔΠ^*N*^ is the same as in the Neo-Hookean case (Eq. (C.14)), and similarly the inequality is calculated as

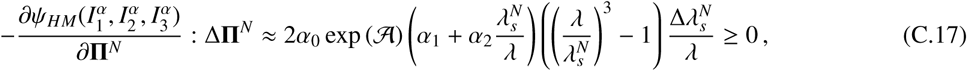

where

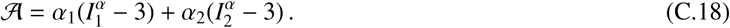

That also indicates since*λ^*N*−1^ < λ^*N*^ <λ* the conditions of the second law are met for sliding.

## Appendix D. Model parameters used in the illustrative examples

The illustrative examples were designed using the noted constitutive relations to demonstrate the important features of the framework with no particular focus on certain tissues; however, in practice, specific constitutive relations should be employed to use the framework for successfully modeling of the inelastic behaviors of tissue. The models generated with this framework may have different number of model parameters. Thus, in general, no specific number of parameters could be associated with the framework. Below we have listed the model parameters used for different examples.

**Table D.1.** Parameters used for comparison of reactive inelasticity to classic models (Fig 6).

| | $\tau$ | $C_1$ | $c$ | $b$ | $(r_0)_s$ |
| --- | --- | --- | --- | --- | --- |
| Formative | 10 | 100 | - | - | - |
| Permanent | $10^{10}$ | 100 | - | - | - |
| Sliding | 0.01 | 100 | 10 | 2 | 1.05 |

**Table D.2.** Parameters used for dynamic response of single bond types (Fig 7).

| | $\tau$ | $C_2$ | $C_3$ | $c$ | $b$ | $(r_0)_s$ | $k$ | $l$ | $(r_0)_D$ |
| --- | --- | --- | --- | --- | --- | --- | --- | --- | --- |
| Formative | 10 | 0.1 | 40 | - | - | - | - | - | - |
| Permanent | $10^{10}$ | 0.1 | 40 | - | - | - | 2 | 1.04 | 1.03 |
| Sliding | 0.01 | 0.1 | 40 | 9 | 2 | 1.01 | - | - | - |

**Table D.3.** Parameters used for the incremental stress relaxation response tests(Fig 8).

| | $\tau$ | $C_2$ | $C_3$ | $E$ | $\nu$ | $\alpha_0$ | $c$ | $b$ | $(r_0)_s$ | $k$ | $l$ | $(r_0)_D$ |
| --- | --- | --- | --- | --- | --- | --- | --- | --- | --- | --- | --- | --- |
| Formative | 10 | 0.1 | 40 | - | - | - | - | - | - | - | - | - |
| Permanent | $10^{10}$ | - | - | 50 | 0.3 | 0.1 | - | - | - | 2 | 1.05 | 1.02 |
| Sliding | 0.01 | - | - | 50 | 0.3 | 0.1 | 8 | 2 | 1.02 | - | - | - |

## References

Alastrué, V., Rodríguez, J., Calvo, B., & Doblaré, M., (2007). Structural damage models for fibrous biological soft tissues. International Journal of Solids and Structures, 44, 5894–5911. URL: http://www.sciencedirect.com/science/article/pii/S0020768307000741http:linkinghub.elsevier.com/retrieve/pii/S0020768307000741. doi:10.1016/j.ijsolstr.2007.02.004.

Andrews, R. D., Tobolsky, A. V., Hanson, E. E. (1946). The Theory of Permanent Set at Elevated Temperatures in Natural and Synthetic Rubber Vulcanizates. Journal of Applied Physics, 17, 352–361. URL: http://aip.scitation.org/doi/10.1063/1.1707724. doi:10.1063/1. 1707724.

Ateshian, G.A. (2015).Viscoelasticity using reactive constrained solid mixtures.Journal of Biomechanics, 48,941–947 URL: http://www.sciencedirect.com/science/article/pii/S0021929015000986. http://linkinghub.elsevier.com/retrieve/pii/S0021929015000986 doi:10.1016/j.jbiomech.2015.02.019.

Caro-Bretelle, A., Gountsop, P., Ienny, P., Leger, R., Corn, S., Bazin, I., & Bretelle, F. (2015). Effect of sample preservation on stress softening and permanent set of porcine skin. Journal of Biomechanics, 48, 3135–3141. doi:10.1016/j.jbiomech.2015.07.014.

Chu, B. M., & Blatz, P. J. (1972). Cumulative microdamage model to describe the hysteresis of living tissue. Annals of Biomedical Engineering, 1, 204–211. URL:http://link.springer.com/10.1007/BF02584207. doi:10.1007/BF02584207.

Coleman, B. D., & Gurtin, M. E. (1967). Thermodynamics with Internal State Variables. The Journal of Chemical Physics, 47, 597–613. URL: http://scitation.aip.org/content/aip/journal/jcp/47/2/10.1063/1.1711937. doi:10.1063/1.1711937.

Connizzo, B. K., & Grodzinsky, A. J. (2017). Tendon exhibits complex poroelastic behavior at the nanoscale as revealed by high-frequency AFM-based rheology. Journal of Biomechanics, 54, 11–18. URL:http://linkinghub.elsevier.com/retrieve/pii/S0021929017300428. doi:10.1016/j.jbiomech.2017.01.029.

Cortes, D. H., & Elliott, D. M. (2014). Accurate prediction of stress in fibers with distributed orientations using generalized high-order structure tensors. Mechanics of Materials, 75, 73–83. URL: http://linkinghub.elsevier.com/retrieve/pii/S0167663614000714. doi:10.1016/j.mechmat.2014.04.006.

Demirkoparan, H., Pence, T. J., & Wineman, A. (2009). On dissolution and reassembly of filamentary reinforcing networks in hyperelastic materials. Proceedings of the Royal Society A: Mathematical, Physical and Engineering Sciences, 465, 867–894. URL: http://rspa.royalsocietypublishing.org/cgi/doi/10.1098/rspa.2008.0360. doi:10.1098/rspa.2008.0360.

Depalle, B., Qin, Z., Shefelbine, S. J., & Buehler, M. J. (2015). Influence of cross-link structure, density and mechanical properties in the mesoscale deformation mechanisms of collagen fibrils. Journal of the Mechanical Behavior of Biomedical Materials, 52, 1–13. doi:10.1016/j.jmbbm.2014.07.008.

Diani, J., Fayolle, B., & Gilormini, P. (2009). A review on the Mullins effect. URL:https://www-sciencedirect-com.udel.idm.oclc.org/science/article/pii/S0014305708006332. doi:10.1016/j.eurpolymj.2008.11.017.

Dittmar, R., Van Rijsbergen, M. M., & Ito, K. (2016). Moderately Degenerated Human Intervertebral Disks Exhibit a Less Geometrically Specific Collagen Fiber Orientation Distribution. Global Spine Journal, 6, 439–446. URL:http://www.ncbi.nlm.nih.gov/pubmed/27433427. doi:10.1055/s-0035-1564805.

Dorfmann, A., & Ogden, R. (2004). A constitutive model for the Mullins effect with permanent set in particle-reinforced rubber. International Journal of Solids and Structures, 41, 1855–1878. URL:http://linkinghub.elsevier.com/retrieve/pii/S0020768303006504. doi:10.1016/j.ijsolstr.2003.11.014.

Eyre, D. R., Weis, M. A., & Wu, J.-J. (2008). Advances in collagen cross-link analysis. Methods, 45, 65–74. doi:10.1016/j.ymeth.2008.01.002.

Fereidoonnezhad, B., Naghdabadi, R., & Holzapfel, G. (2016). Stress softening and permanent deformation in human aortas: Continuum and computational modeling with application to arterial clamping. Journal of the Mechanical Behavior of Biomedical Materials, 61, 600–616. URL: http://linkinghub.elsevier.com/retrieve/pii/S1751616116300467. doi:10.1016/j.jmbbm.2016.03.026.

Fung, Y.-C. (1993). Biomechanics. New York, NY: Springer New York. URL: http://link.springer.com/10.1007/978-1-4757-2257-4.doi:10.1007/978-1-4757-2257-4.

Green, M. S., & Tobolsky, A. V. (1946). A New Approach to the Theory of Relaxing Polymeric Media. The Journal of Chemical Physics, 14, 80–92. URL: http://aip.scitation.org/doi/10.1063/1.1724109. doi:10.1063/1.1724109.

Herod, T. W., Chambers, N. C., & Veres, S. P. (2016). Collagen fibrils in functionally distinct tendons have di ering structural responses to tendon rupture and fatigue loading. Acta Biomaterialia, 42, 296–307. URL: http://linkinghub.elsevier.com/retrieve/pii/S174270611630294X. doi:10.1016/j.actbio.2016.06.017.

Holmes, M. H., & Mow, V. C. (1990). The nonlinear characteristics of soft gels and hydrated connective tissues in ultrafiltration. Journal of biomechanics, 23, 1145–56. URL: http://www.ncbi.nlm.nih.gov/pubmed/2277049.

Holzapfel, G. (2000). Nonlinear Solid Mechanics: A Continuum Approach for Engineering. URL: http://www.wiley.com/WileyCDA/WileyTitle/productCd-0471823198.html.

Holzapfel, G. A., & Simo, J. C. (1996). A new viscoelastic constitutive model for continuous media at finite thermomechanical changes. International Journal of Solids and Structures, 33, 3019–3034. URL: http://linkinghub.elsevier.com/retrieve/pii/00207 http://linkinghub.elsevier.com/retrieve/pii/0020768395002634. doi:10.1016/0020-7683(95)00263-4.

Horstemeyer, M. F., & Bammann, D. J. (2010).Historical review of internal state variable theory for inelasticity. International Journal of Plasticity, 26, 1310–1334. URL: http://linkinghub.elsevier.com/retrieve/pii/S0749641910000847. doi:10.1016/j.ijplas.2010.06.005.

Huang, C. Y., Mow, V. C., & Ateshian, G. A. (2001). The role of flow-independent viscoelasticity in the biphasic tensile and compressive responses of articular cartilage. Journal of biomechanical engineering, 123, 410–417. URL: http://biomechanical.asmedigitalcollection.asme.org/data/Journals/JBENDY/26190/410{_}1.pdf. doi:10.1115/1.1392316.

Jacobs, N. T., Cortes, D. H., Peloquin, J. M., Vresilovic, E. J., & Elliott, D. M. (2014). Validation and application of an intervertebral disc finite element model utilizing independently constructed tissue-level constitutive formulations that are nonlinear, anisotropic, and time-dependent. Journal of Biomechanics, 47, 2540–2546. URL: http://linkinghub.elsevier.com/retrieve/pii/S0021929014003479. doi:10.1016/j.jbiomech.2014.06.008.

Ju, J. W. (1990). Isotropic and Anisotropic Damage Variables in Continuum Damage Mechanics. Journal of Engineering Mechanics, 116, 2764–2770. doi:10.1061/ASCE0733-9399(1990)116:12(2764).

Kachanov, L. M. (1968). Introduction to continuum damage mechanics. Springer Science & Business Media.

Khan, A. S., & Huang, S. (1995). Continuum theory of plasticity. John Wiley & Sons.

Krajcinovic, D. (2000). Damage mechanics: accomplishments, trends and needs. International Journal of Solids and Structures, 37, 267–277. URL: http://www.sciencedirect.com/science/article/pii/S0020768399000815. doi:10.1016/S0020-7683(99)00081-5.

Lee, A. H., Szczesny, S. E., Santare, M. H., & Elliott, D. M. (2017). Investigating mechanisms of tendon damage by measuring multi-scale recovery following tensile loading. Acta Biomaterialia, 57, 363–372. URL: http://www.sciencedirect.com/science/article/pii/S1742706117302386 http://linkinghub.elsevier.com/retrieve/pii/S1742706117302386. doi:10.1016/j.actbio.2017.04.011.

Lee, E. H. (1969). Elastic-Plastic Deformation at Finite Strains. Journal of Applied Mechanics, 36, 1. URL: http://appliedmechanics.asmedigitalcollection.asme.org/article.aspx?articleid=1398837. doi:10.1115/1.3564580.

Lemaitre, J. (1984). How to use damage mechanics. Nuclear Engineering and Design, 80, 233–245. URL:http://linkinghub.elsevier.com/retrieve/pii/0029549384901699. doi:10.1016/0029-5493(84)90169-9.

Li, W. (2016). Damage Models for Soft Tissues: A Survey. Journal of Medical and Biological Engineering, 36, 285–307. URL: http://link.springer.com/10.1007/s40846-016-0132-1". doi:10.1007/s40846-016-0132-1.

Li, Y., & Yu, S. M. (2013). Targeting and mimicking collagens via triple helical peptide assembly. Current Opinion in Chemical Biology, 17, 968–975. URL: http://www.ncbi.nlm.nih.gov/pubmed/24210894 http://www.pubmedcentral.nih.gov/articlerender.fcgi?artid=PMC3863647http://linkinghub.elsevier.com/retrieve/pii/S1367593113001841. doi:10.1016/j.cbpa.2013.10.018.

Lubarda, V. A. (2004). Constitutive theories based on the multiplicative decomposition of deformation gradient: Thermoelasticity, elastoplasticity, and biomechanics. Applied Mechanics Reviews, 57, 95. URL: http://newmaeweb.ucsd.edu/{~}vlubarda/research/pdfpapers/amr-04.pdf http://appliedmechanicsreviews.asmedigitalcollection.asme.org/article.aspx?articleid=1397944. doi:10.1115/1.1591000.arXiv:4644220693.

Lynch, H. A., Johannessen, W., Wu, J. P., Jawa, A., & Elliott, D. M. (2003). Effect of fiber orientation and strain rate on the nonlinear uniaxial tensile material properties of tendon. Journal of biomechanical engineering, 125, 726–731. doi:10.1115/1.1614819.

Maher, E., Creane, A., Lally, C., & Kelly, D. J. (2012). An anisotropic inelastic constitutive model to describe stress softening and permanent deformation in arterial tissue. Journal of the Mechanical Behavior of Biomedical Materials, 12, 9–19. doi:10.1016/j.jmbbm.2012.03.001.

Meng, F., Pritchard, R. H., & Terentjev, E. M. (2016). Stress Relaxation, Dynamics, and Plasticity of Transient Polymer Networks. Macromolecules, 49, 2843–2852. URL:http://pubs.acs.org/doi/abs/10.1021/acs.macromol.5b02667. doi:10.1021/acs.macromol.5b02667.

Meng, F., & Terentjev, E. (2016). Transient Network at Large Deformations: Elastic–Plastic Transition and Necking Instability. Polymers, 8, 108.URL: http://www.mdpi.com/2073-4360/8/4/108. doi:10.3390/polym8040108.

Muliana, A., Rajagopal, K. R., Tscharnuter, D., & Pinter, G. (2016). A nonlinear viscoelastic constitutive model for polymeric solids based on multiple natural configuration theory. International Journal of Solids and Structures, 100-101, 95–110. URL:http://linkinghub.elsevier.com/retrieve/pii/S0020768316301780. doi:10.1016/j.ijsolstr.2016.07.017.

Mullins, L. (1969). Softening of Rubber by Deformation. Rubber Chemistry and Technology, 42, 339–362. URL:http://rubberchemtechnol.org/doi/abs/10.5254/1.3539210. doi:10.5254/1.3539210.

Murakami, S. (2012). Continuum Damage Mechanics volume 185 of Solid Mechanics and Its Applications. Dordrecht: Springer Netherlands. URL:http://link.springer.com/10.1007/978-94-007-2666-6. doi:10.1007/978-94-007-2666-6.

Naghdi, P., & Trapp, J. (1975). The significance of formulating plasticity theory with reference to loading surfaces in strain space. International Journal of Engineering Science, 13, 785–797. URL:http://linkinghub.elsevier.com/retrieve/pii/0020722575900804. doi:10.1016/0020-7225(75)90080-4.

Natali, A., Pavan, P., Carniel, E., Lucisano, M., & Taglialavoro, G. (2005). Anisotropic elasto-damage constitutive model for the biomechanical analysis of tendons. Medical Engineering & Physics, 27, 209–214. URL:http://linkinghub.elsevier.com/retrieve/pii/S1350453304002036. doi:10.1016/j.medengphy.2004.10.011.

Nims, R. J., & Ateshian, G. A. (2017). Reactive Constrained Mixtures for Modeling the Solid Matrix of Biological Tissues. Journal of Elasticity, URL:http://linkspringer.com/10.1007/s10659-017-9630-9. doi:10.1007/s10659-017-9630-9.

Nims, R. J., Durney, K. M., Cigan, A. D., Dussèaux, A., Hung, C. T., & Ateshian, G. A. (2016). Continuum theory of fibrous tissue damage mechanics using bond kinetics: application to cartilage tissue engineering. Interface Focus, 6, 20150063. URL:http://rsfs.royalsocietypublishing.org/lookup/doi/10.1098/rsfs.2015.0063. doi:10.1098/rsfs.2015.0063.

Peña, E. (2011). Prediction of the softening and damage effects with permanent set in fibrous biological materials. Journal of the Mechanics and Physics of Solids, 59, 1808–1822. URL:http://linkinghub.elsevier.com/retrieve/pii/S0022509611001220. doi:10.1016/ j.jmps.2011.05.013.

Peña, E. (2014). Computational aspects of the numerical modelling of softening, damage and permanent set in soft biological tissues. Computers & Structures, 130, 57–72. URL: http://www.sciencedirect.com/science/article/pii/S0045794913002642. doi:10.1016/j.compstruc.2013.10.002.

Rajagopal, K., & Wineman, A. (1992). A constitutive equation for nonlinear solids which undergo deformation induced microstructural changes. International Journal of Plasticity, 8, 385–395. URL: http://www.linkinghub.elsevier.com/retrieve/pii/074964199290056I. doi:10.1016/0749-6419(92)90056-I.

Rajagopal, K. R., & Srinivasa, A. R. (2004). On the thermo mechanics of materials that have multiple natural configurations Part I: Viscoelasticity and classical plasticity. URL: https://link.springer.com/content/pdf/10.1007/s00033-004-4020-0.pdf. doi:10.1007/s00033-004-4019-6.

Reese, S., & Govindjee, S. (1998). A theory of finite viscoelasticity and numerical aspects. International Journal of Solids and Structures, 35, 3455–3482. URL: http://linkinghub.elsevier.com/retrieve/pii/S0020768397002175. doi:10.1016/S0020-7683(97)00217-5.

Reiser, K., McCormick, R. J., & Rucker, R. B. (1992). Enzymatic and nonenzymatic cross-linking of collagen and elastin. FASEB journal: official publication of the Federation of American Societies for Experimental Biology, 6, 2439–49. URL: http://www.ncbi.nlm.nih.gov/pubmed/1348714.

Safa, B. N. (2018). ReactiveBond. URL: https://github.com/BabakNSafa/ReactiveBond/.

Schmidt, T., Balzani, D., & Holzapfel, G. (2014). Statistical approach for a continuum description of damage evolution in soft collagenous tissues. Computer Methods in Applied Mechanics and Engineering, 278, 41–61. URL: http://www.sciencedirect.com/science/article/pii/S0045782514001364. doi:10.1016/j.cma.2014.04.011.

Scott, K. W., & Stein, R. S. (1953). A Molecular Theory of Stress Relaxation in Polymeric Media. The Journal of Chemical Physics, 21, 1281–1286. URL: http://aip.scitation.org/doi/10.1063/1.1699181. doi:10.1063/1.1699181.

Shen, Z. L., Dodge, M. R., Kahn, H., Ballarini, R., & Eppell, S. J. (2008). Stress-Strain Experiments on Individual Collagen Fibrils. Biophysical Journal, 95, 3956–3963. URL: http://linkinghub.elsevier.com/retrieve/pii/S0006349508785345. doi:10.1529/biophysj. 107.124602.

Simo, J. (1987). On a fully three-dimensional finite-strain viscoelastic damage model: Formulation and computational aspects. Computer Methods in Applied Mechanics and Engineering, 60, 153–173. URL: http://linkinghub.elsevier.com/retrieve/pii/0045782587901071. doi:10.1016/0045-7825(87)90107-1.

Simo, J. (1988). A framework for finite strain elastoplasticity based on maximum plastic dissipation and the multiplicative decomposition: Part I. Continuum formulation. Computer Methods in Applied Mechanics and Engineering, 66, 199–219. doi:10.1016/0045-7825(88)90076-X.

Simo, J. C., & Hughes, T. J. R. (1998). Computational Inelasticity. Interdisciplinary Applied Mathematics. New York: Springer-Verlag. URL: http://link.springer.com/10.1007/b98904. doi:10.1007/b98904.

Spencer, A. J. (1984). Continuum Theory of the Mechanics of Fibre-Reinforced Composites. Vienna: Springer Vienna. URL: http://link.springer.com/10.1007/978-3-7091-4336–0. doi:10.1007/978-3-7091-4336-0.

Szczesny, S. E., & Elliott, D. M. (2014). Interfibrillar shear stress is the loading mechanism of collagen fibrils in tendon. Acta Biomaterialia, 10, 2582–2590. URL: http://www.ncbi.nlm.nih.gov/pubmed/24530560. doi:10.1016/j.actbio.2014.01.032.

Tanaka, F., & Edwards, S. (1992). Viscoelastic properties of physically crosslinked networks. Journal of Non-Newtonian Fluid Mechanics, 43, 273–288. URL: http://linkinghub.elsevier.com/retrieve/pii/037702579280028V. doi:10.1016/0377-0257(92)80028-V.

Tobolsky, A. V., & Andrews, R. D. (1945). Systems Manifesting Superposed Elastic and Viscous Behavior. The Journal of Chemical Physics, 13, 3–27. URL: http://aip.scitation.org/doi/10.1063/1.1723966. doi:10.1063/1.1723966.

Veres, S. P., Harrison, J. M., & Lee, J. M. (2014). Mechanically overloading collagen fibrils uncoils collagen molecules, placing them in a stable, denatured state. Matrix Biology,. doi:10.1016/j.matbio.2013.07.003.

Von Forell, G. A., & Bowden, A. E. (2014). A damage model for the percutaneous triple hemisection technique for tendo-achilles lengthening. Journal of Biomechanics, 47, 3354–3360. URL: http://www.sciencedirect.com/science/article/pii/S0021929014004308http://linkinghub.elsevier.com/retrieve/pii/S0021929014004308. doi:10.1016/j.jbiomech.2014.08.006.

Weisbecker, H., Pierce, D. M., Regitnig, P., & Holzapfel, G. A. (2012). Layer-specific damage experiments and modeling of human thoracic and abdominal aortas with non-atherosclerotic intimal thickening. Journal of the Mechanical Behavior of Biomedical Materials, 12, 93–106. URL: www.sciencedirect.comwww.elsevier.com/locate/jmbbm. doi:10.1016/j.jmbbm.2012.03.012.

Wineman, A. (2009). On the mechanics of elastomers undergoing scission and cross-linking. International Journal of Advances in Engineering Sciences and Applied Mathematics, 1, 123–131. URL: http://link.springer.com/10.1007/s12572-010-0004-9. doi:10.1007/ s12572-010-0004-9.

Woo, S. L.-Y., Simon, B. R., Kuei, S. C., & Akeson, W. H. (1980). Quasi-Linear Viscoelastic Properties of Normal Articular Cartilage. Journal of Biomechanical Engineering, 102, 85. URL: http://biomechanical.asmedigitalcollection.asme.org/article.aspx?articleid=1394855. doi:10.1115/1.3138220.

Zhang, W., & Sacks, M. S. (2017). Modeling the response of exogenously crosslinked tissue to cyclic loading: The effects of permanent set. Journal of the Mechanical Behavior of Biomedical Materials, 75, 336–350. URL: http://linkinghub.elsevier.com/retrieve/pii/S1751616117302990. doi:10.1016/j.jmbbm.2017.07.013.

Zitnay, J. L., Li, Y., Qin, Z., San, B. H., Depalle, B., Reese, S. P., Buehler, M. J., Yu, S. M., & Weiss, J. A. (2017). Molecular level detection and localization of mechanical damage in collagen enabled by collagen hybridizing peptides. Nature Communications, 8, 14913. URL: http://www.nature.com/doifinder/10.1038/ncomms14913. doi:10.1038/ncomms14913.

